# Lessons learned from a Kaggle challenge for particle picking in cryo-electron tomography

**DOI:** 10.1101/2025.11.03.686153

**Authors:** Ariana Peck, Joshua Hutchings, Jonathan Schwartz, Yue Yu, Utz H. Ermel, Saugat Kandel, Dari Kimanius, Zhuowen Zhao, Shawn Zheng, Brendan Artley, David List, Sergio A. Silva, Walter Reade, Jeremy Asuncion, Kira Evans, Jessica Gadling, Kandarp Khandwala, Suzette McCanny, Dannielle G. McCarthy, Jun Xi Ni, Janeece Pourroy, Manasa Venkatakrishnan, Zun Shi Wang, David A. Agard, Clinton S. Potter, Bridget Carragher, Kyle Harrington, Mohammadreza Paraan

## Abstract

The difficulty of particle picking in cryo-electron tomography remains a barrier to routine in situ structure determination. Machine learning is well-suited to overcome this bottleneck with efficient algorithms that generalize across molecular species. To spur new algorithm development, we held a three-month Kaggle challenge that tasked contestants with annotating five molecular species across hundreds of experimental tomograms. This competition successfully engaged >1000 participants from diverse fields and delivered particle pickers that outperformed existing state-of-the-art. Systematic comparisons of the contestants’ submissions revealed the tolerance of subtomogram averaging to moderate but not severe over-picking and underscored the need for more robust measures of annotation quality. The winning models also highlighted the importance of data augmentation to overcome limited training data. All competition tomograms along with the ground truth and winning teams’ annotations have been released on the CryoET Data Portal as a resource to benchmark current and future tools for particle picking.

## Introduction

Cryo-electron tomography (cryoET) is a powerful technique for visualizing cellular ultrastructure and molecular complexes in situ^1^. In this imaging modality, a vitrified specimen is illuminated by an electron beam and systematically tilted while the corresponding projection images are collected. The resulting tilt-series is reconstructed into a tomogram that captures the biological state with molecular detail at the moment of freezing. Identifying sufficient copies of a target molecular species in the crowded environment of the cell enables subtomogram averaging (STA), a method to align and average these individual copies in order to resolve the molecule’s structure^2^. STA can achieve subnanometer resolution^3,4^ and reveal conformational ensembles^5^ but generally requires annotating tens of thousands of molecular copies across tens to hundreds of tomograms.

Molecular annotation remains the principal bottleneck of cryoET data processing because time-consuming manual effort is often needed to obtain the large and highly curated set of particles required for high-resolution STA. Annotation, which is also referred to as particle picking or labeling, is made difficult by multiple factors, including the data’s extremely low signal-to-noise ratio (SNR), the missing wedge of information that blurs the signal along the imaging axis^6^, the extreme molecular crowding of cellular tomograms, and the diversity and heterogeneity of the species of interest. The conventional annotation method of template matching is computationally expensive and depends on having a suitable template for the target of interest^6–9^. As a result, this technique is restricted to molecular species with known structures and generalizes poorly to conformationally heterogeneous targets and isolated complexes of low molecular weight^10,11^. Machine learning (ML) algorithms are well-suited to provide more scalable and versatile solutions to this annotation bottleneck. Despite the recent development of numerous ML particle picking algorithms^12–18^, the fact that the field has not coalesced behind a single method suggests that advances are still needed to meet the diverse needs of the cryoET community.

Community challenges have proven invaluable for catalyzing new methods development and validating existing tools in structural biology. Previously the field held two consecutive annotation challenges that benchmarked several ML algorithms developed by cryoET experts against a synthetic dataset consisting of ten tomograms^19–21^. Taking inspiration from that precedent and notable community challenges in other domains of structural biology^22,23^, we sought to take the next critical step by launching a challenge of considerably expanded scope. We based our challenge on a large experimental dataset spanning nearly 500 annotated tomograms and five scored targets of different sizes and shapes to favor efficient solutions robust to experimental artifacts and capable of generalizing across species^24^. We launched our challenge on Kaggle to engage this platform’s broad ML community and tap into expertise beyond the cryoET field. This three-month competition yielded a comprehensive comparison of particle picking algorithms that spanned ∼28,000 solutions from 1131 contestants across 76 countries. Collectively, these results highlighted strategies that enabled models to perform robustly under the realistic constraint of limited training data and offered insights into how improvements in model selectivity impact STA. The annotations from the winning submissions, which outperformed our benchmark solution^24^ and in-field particle picking algorithms applied out-of-the-box^13,17^, have all been released on the CryoET Data Portal^25^ alongside the full dataset to enable validating new and existing tools against the results of this community challenge.

## Results

### Competition Format

Our competition was centered on a phantom dataset designed to benchmark multi-species annotation algorithms^24^. This experimental dataset spanned 492 high-quality tomograms and ∼60k ground truth labels across six particle classes (Fig. 1a). Five of these particles — apoferritin, beta-amylase, beta-galactosidase, thyroglobulin, and virus-like particles — were mixed with cellular lysate enriched for lysosomes. These organelles enforced a sample thickness of ∼200 nm similar to most lamella samples and provided space for layers of particles to be stacked along the tomogram depth, making annotation more challenging than in single-particle cryoEM data. The enriched lysate contained ribosomes, the sixth target particle, in abundance, and numerous other molecular species that served as natural decoys to confound annotation algorithms. Collectively, these six target particles spanned a range of molecular weights (268 kDa - 4.3 MDa) and distinct shapes to encourage algorithms that generalize to diverse species. The ground truth annotations were meticulously curated to the best of the abilities of a team of cryoET experts, and their high quality was validated by STA, which yielded refined maps at resolutions of 3.6-11.5 Å^24^. However, these ground truth labels are necessarily imperfect given the experimental nature of the data^24^.

**Fig. 1:**
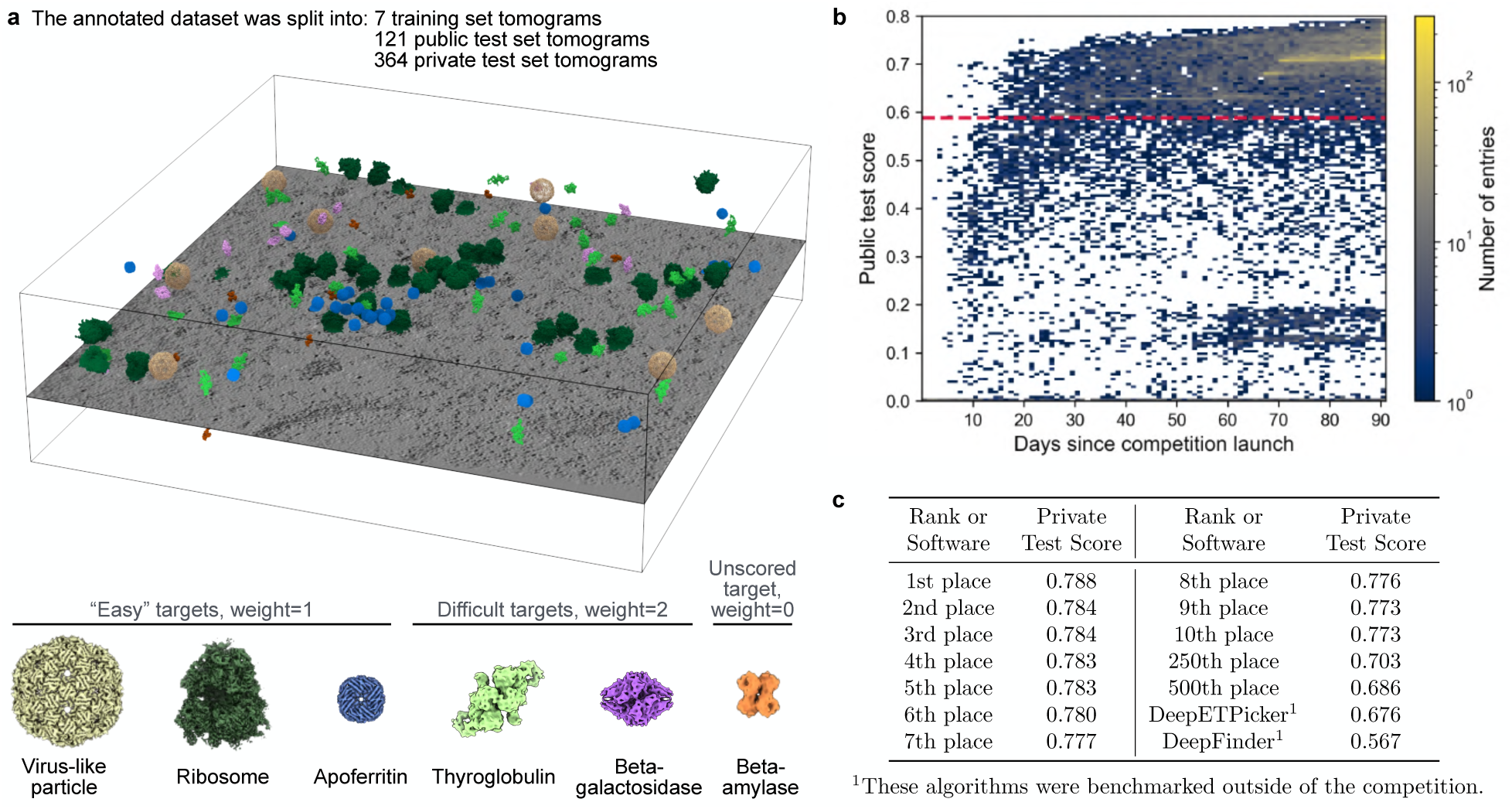
Overview of the competition. **a.** The phantom dataset^24^ consisted of 492 annotated tomograms that were split into a training set (7 tomograms), a public test set (121 tomograms), and a private test set (364 tomograms). (upper) A slice through a representative tomogram from the training set and annotated particles throughout the volume are visualized. (lower) The six annotation targets and the class weights applied during scoring are shown. The particles are drawn to scale. **b.** The evolution of the public test scores is tracked over the course of the competition for all submissions. The dashed red line indicates the public test score achieved by our benchmark solution^24^, which was adapted from DeepFinder^13^. **c.** The private test scores achieved by the top 10 contestants, two lower-ranked submissions, and two existing particle picking algorithms benchmarked outside of the competition are listed.

For the competition, this phantom dataset was split into three subsets. The first subset was the training set, which consisted of only seven annotated tomograms to reflect the amount of data that is reasonable for a cryoET expert to annotate by hand. The second subset was the public test set, which comprised 25% of the remaining dataset. These 121 annotated tomograms were withheld throughout the competition, but contestants were provided with a model score against these test data for each solution they submitted. Many contestants reported that these public leaderboard scores were useful for guiding model design and often proved more reliable than cross-validation due to the small training set size^26^. The third and final subset was the private test set, which consisted of the remaining 364 tomograms. In this case, both the annotated tomograms and the model scores were withheld since these scores were used to rank the submissions and select the winning teams.

Submissions were evaluated based on the F_®_ score. This generalization of the F_1_ score includes a β parameter that tunes the relative penalties applied to false negatives and false positives. For our competition, we set β=4 to prioritize recall over precision. Our rationale is that we wanted to severely penalize models with poor recall given the high quality of the ground truth labels. However, we could not guarantee that we had not missed any bona fide particles during our annotation process, so we made scoring lenient towards contestants’ false positives in case any corresponded to valid particles. Further, whereas false positives can be filtered out by curation, false negatives cannot be recovered after picking, a difference that favors algorithms that over-rather than under-pick. The F_β_ score was then calculated for each particle class across the test tomograms, and an aggregate score was computed from the weighted average of the per-particle F_β_ scores. We assigned a weight of 1 to the particle classes (virus-like particles, ribosomes, and apoferritin) we considered easy to annotate and a weight of 2 to the challenging species (thyroglobulin and beta-galactosidase) to favor algorithms that performed robustly for difficult targets. Ultimately we decided against scoring beta-amylase due to low confidence in the quality of our annotations for this particle class. However, ground truth labels for beta-amylase were provided for the training data.

### Statistical analysis of the submissions

On November 6, 2024, we launched our competition on Kaggle to tap into this platform’s active community of ML experts. During the three months of the competition, we attracted 1131 participants from across 76 countries who collectively submitted nearly 28,000 solutions. Kaggle’s “play to learn” culture of competitive collaboration was evident in the animated discussions on the competition’s forum and the number of contestants who teamed up to merge approaches. These high levels of participation and engagement drove the competition forward. By the end of our challenge, nearly three quarters of submissions outperformed the public test score achieved by our benchmark solution (Fig. 1b). The top-performing models also outperformed two in-field particle picking algorithms, DeepFinder^13^ and DeepETPicker^17^, that were trained on the training set and applied to the test data out-of-the-box without any hyperparameter tuning (Fig. 1c). The rapid improvement in the contestants’ model performance was supported by the significant compute resources that Kaggle provided. The contestants collectively used ∼10,000 GPU-hours donated by Kaggle, with all but one of the winning teams submitting >100 solutions during the competition (Extended Data Fig. 1a). Submissions were subject to Kaggle’s 12-hour runtime limit, which incentivized contestants to balance efficiency and accuracy. Importantly, model performance was consistent between the private and public test data, indicating that the phantom dataset had been successfully split into subsets with similar distributions across particle classes (Extended Data Fig. 1b). This observation also suggested that the contestants were not overfitting to the public test data despite leveraging the public leaderboard scores to guide model design.

Given that the test scores aggregate information across all particle classes, we assessed model performance for each target species. The fraction of ground truth labels recovered (recall) exhibited a few different trends (Fig. 2a). For the virus-like particles, we observed robust recovery of the ground truth labels across a range of test scores, presumably because its size and symmetry make it the easiest target to annotate. However, there was a noticeable fall-off in the fraction of ground truth recovered for the other species as test scores decreased. Curiously, for thyroglobulin, many submissions with intermediate test scores recovered a slightly higher fraction of the ground truth labels than the top-ranked submissions. Despite this pattern, recovery of the ground truth saturated at a lower fraction for the difficult annotation targets than for the easy targets. We also examined the over-picking ratio — i.e., the ratio of the number of annotations a contestant submitted to the ground truth counts — as a rough measure of model selectivity (Fig. 2b). For the virus-like particles and two challenging targets, this metric increased significantly with decreasing test scores, indicating poorer model selectivity. In the case of thyroglobulin, severe over-picking may account for the modest increase in ground truth recovery at intermediate test scores, since indiscriminate labeling would make it statistically more likely to recover ground truth annotations. On the other hand, the over-picking ratio was more stable for the ribosomes and apoferritin, possibly due to their higher abundance compared to the other species. Analysis of inter-class duplicates — i.e., coordinates assigned to multiple particle types — confirmed improved model selectivity with increasing test score (Extended Data Fig. 2).

**Fig. 2:**
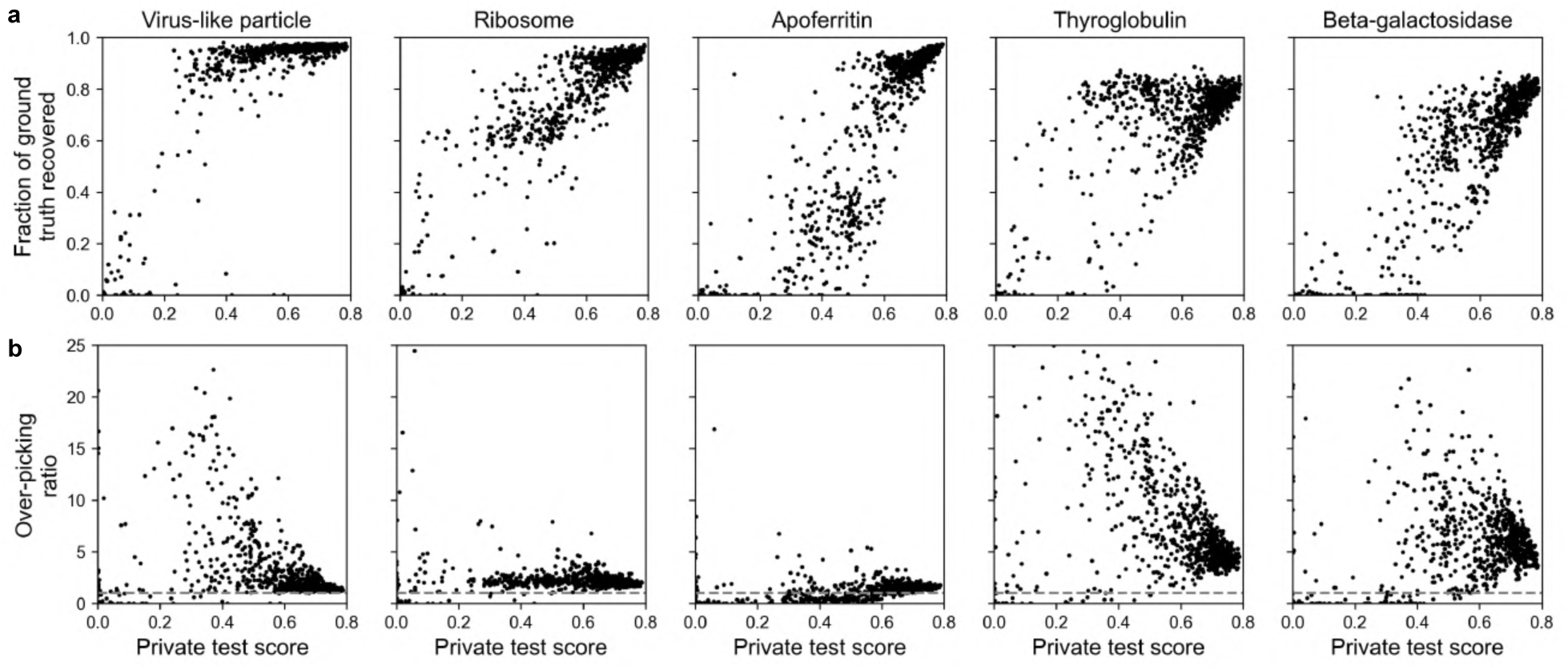
Comparison of contestants’ submissions to the ground truth. **a.** The fraction of ground truth recovered and **b.** the over-picking ratio are plotted as a function of the private test score for the indicated particle class for the ∼24k valid submissions received during the competition. The over-picking ratio refers to the ratio of the number of annotations a contestant submitted compared to the number of ground truth labels for each particle class. The dashed line indicates an over-picking ratio of 1, and the y-axis is capped at an over-picking ratio of 25.

### Structural analysis of select submissions

In cryoET, annotation is often performed with the aim of subtomogram averaging, in which the annotated copies of the molecule are aligned and averaged together to determine a 3D map of the molecule’s structure^2^. Annotation quality can then be assessed based on the map’s resolution, which reflects the quantity and quality (structural coherence) of the picks. To assess how model performance impacts this critical downstream task, we compared the refined maps from the annotations submitted by the 10 winners and 10 lower-ranked teams. The latter were randomly selected as representative submissions that scored between the winning teams’ models and our benchmark solution. Consensus picks from the top 5 teams and the predictions from DeepFinder^13^ and DeepETPicker^17^ were also assessed. To ensure a systematic comparison, all sets of annotations were processed by an automated, reference-based STA pipeline in RELION-5^27^ that generated refined maps from the full set of particles, without any classification, curation, or manual parameter tuning. Contrast transfer function (CTF) refinement and Bayesian polishing, which are often performed iteratively to maximize the high-resolution limit of the reconstruction, were also omitted for computational expediency.

Even without applying the curation steps that are routine during STA, nearly all Kaggle teams and both in-field algorithms yielded maps that were comparable in quality to the ground truth maps for the easy annotation targets (Extended Data Table 1, Extended Data Figs. 3-4). By contrast, map resolution for the difficult targets provided a better diagnostic of model performance, with the lower-ranked teams and DeepFinder often yielding artifactual reconstructions (Extended Data. Fig. 5, Supplemental Note 1). These patterns were reflected in the trends between map resolution and statistical measures of model performance (Fig. 3a). Map quality tended to deteriorate with decreasing test score for the challenging targets, consistent with the higher weights applied to these species. Although no correlation was observed between the fraction of ground truth recovered and resolution, the lowest-quality maps were all characterized by extreme over-picking. The threshold at which over-picking was problematic was species-dependent, presumably because STA’s tolerance to false positives depends on the particle’s size and structural features.

**Fig. 3:**
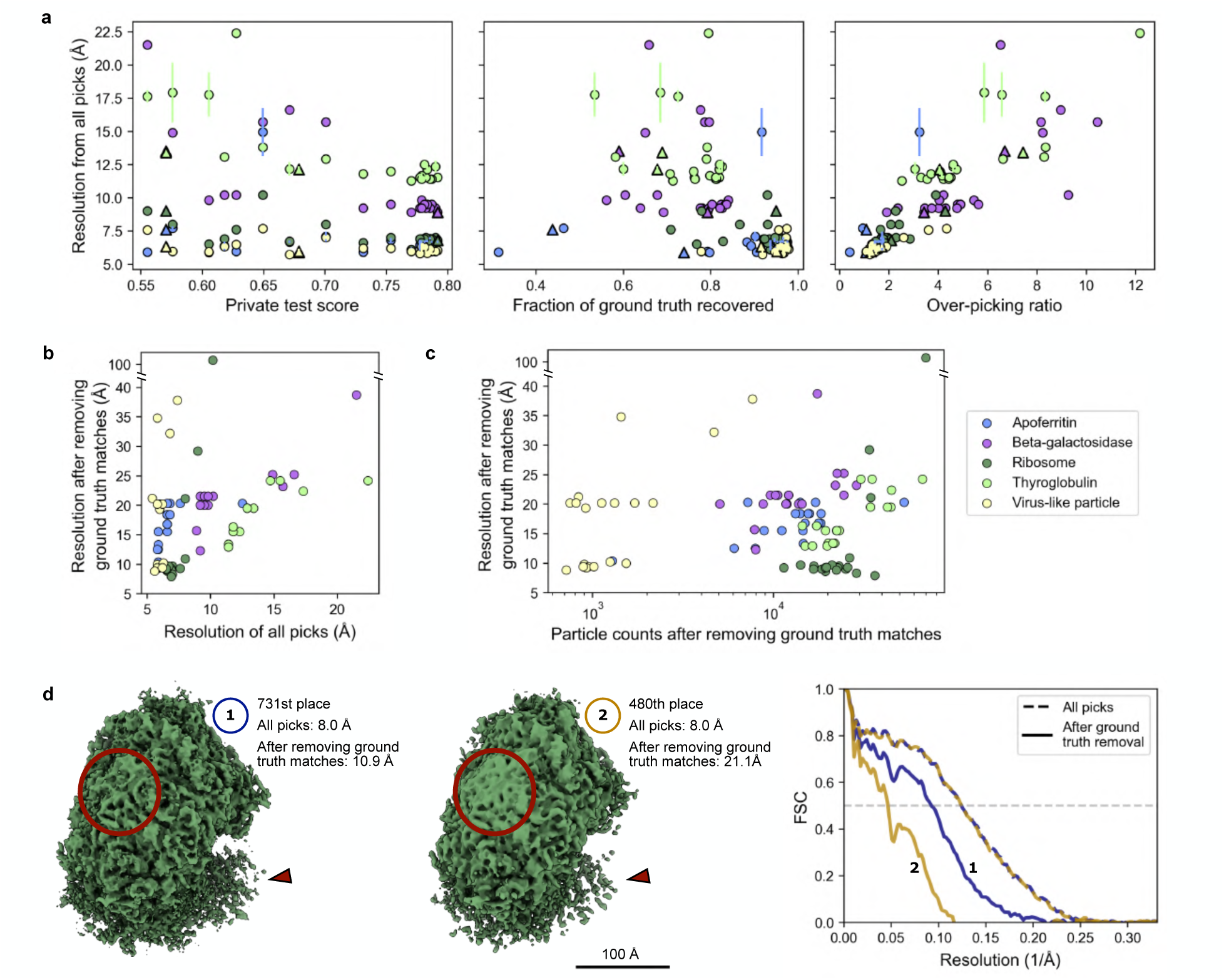
Map quality is sensitive to extreme over-picking but tolerates a moderate degree of contamination. **a.** Resolution estimates for the refined maps from the annotations of 20 Kaggle teams, the top 5 teams’ consensus picks, and two in-field methods are plotted as a function of the private test score (left), the fraction of ground truth recovered (center), and the over-picking ratio (right). STA for the apoferritin, thyroglobulin, and virus-like particle classes was run in triplicate (Supplemental Note 2); for these species, the mean resolution and standard deviation from the three independent runs are plotted. The datapoints corresponding to the in-field methods are plotted as triangles. **b-d.** The picks that matched the ground truth labels were then removed from each of the Kaggle team’s annotations, and STA was performed on the remaining picks. The resolution of the resulting maps after ground truth removal is shown as a function of **b.** the resolution of the corresponding complete sets of picks and **c.** particle counts. **d.** An example case is shown in which the picks from two teams initially yielded maps at the same resolution but at very different resolutions after the ground truth matches were removed. (left) Loss of secondary structural features and diminished density for the small subunit are indicated in red. (The latter does not impact the resolution estimate since the small subunit was masked out for the FSC calculation). (right) The FSC curves are plotted for the maps before (dashed lines) and after (solid lines) omitting the ground truth matches.

One limitation of our scoring metric is that it penalizes all false positives equally, despite the expectation that these contain valid particles missing from the ground truth labels in addition to contamination. To evaluate these presumed false positives, we applied our automated STA pipeline to the teams’ annotations after discarding picks that matched the ground truth labels. Even with the true positives omitted, several teams recovered the correct overall structure for each species (Fig. 3b, Extended Data Fig. 6a). The exception was apoferritin, despite valid particles dominating the consensus false positives (Extended Data Fig. 7a). These results justified the leniency set by our F_β_ score and suggested that many models outperformed our best-effort annotation process that generated the ground truth labels. The variability in map resolution from the teams’ presumed false positives also indicated significant differences in annotation accuracy between teams. However, this accuracy in general could not be predicted from the map quality of the complete set of annotations (Fig. 3b). In multiple cases, sets of picks that produced maps at the same resolution yielded maps that differed by >10 Å after the ground truth matches were removed (Fig. 3d, Extended Data Fig. 6b-c), and these discrepancies could not be accounted for by particle numbers (Fig. 3c). These results underscore the limited sensitivity of STA to a moderate number of false positives since sufficient structural coherence in a set of picks can drive refinement to the correct consensus structure and mask the presence of contamination. The time and compute resources required by STA made it prohibitive for scoring the competition, but more robust measures of annotation quality are needed to quantify differences in model performance.

### Common themes of the winning models

The consistently high performance of the top-ranked submissions raised the question of what made the underlying models so effective. In terms of architecture, the 10 winning models all relied on 3D U-Nets as the backbone, often connected to 2D and/or 3D convolutional neural network (CNN) encoders (Table 1, see Extended Data Table 2 for links to the winners’ reports). The exceptions were the 1st and 2nd place teams, which respectively implemented a YOLO-like object detection model and leveraged residual neural network (ResNet) backbones in combination with 3D U-Nets. Though the 10th place winner wryly referred to 3D U-Nets as a “vintage” architecture, this appeared to be the most effective tool for this challenge. As the winning team noted, more modern architectures like vision transformers dedicate significant representational capacity to modeling global contextual information that was not needed to accurately assign localized regions of the phantom tomograms to specific particle classes. Whether additional context would benefit in situ annotation remains an open question. Further, the number of parameters typical of vision transformers and even large U-Nets made them susceptible of over-fitting to the limited training data. As a result, the winning models all converged on lightweight 3D U-Nets, though there was variability in the choice of loss function and whether contestants found pre-trained weights from either foundation models or synthetic data beneficial (Table 1).

**Table 1:** Overview of the winning models. General features of the models from the top 10 teams are compared. ^a^Some teams used CNN-style encoders to first transform the imaging data into embeddings before feeding them to the backbone models, while others fed the imaging data directly into the backbone. Across the top 10 teams, the backbone architectures were predominantly 3D U-Nets. ^b^Most teams ensembled multiple models with varying numbers of parameters. ^c^CE: cross entropy, IOU: intersection over union, MSE: mean square error. The models are described in depth in the winners’ reports (Extended Data Table 2).

| Rank | Architecture<sup>a</sup> |  | Parameters per model<sup>b</sup> | Number of models | Pretrained weights | Loss function<sup>c</sup> |
| --- | --- | --- | --- | --- | --- | --- |
|  | Encoder | Backbone |  |  |  |  |
| 1 | 3D CNN | 3D U-Net | 71M | 12 | No | CE |
|  | N/A | YOLO-like | 19 - 31M | 15 | Yes | Varifocal, IOU |
| 2 | 2D / 3D CNN | 3D U-Net | 870K - 14M | 10 | No | Tversky, Dice, CE |
| 3 | 3D CNN | 3D U-Net | 11 - 131M | 4 | No | CE |
| 4 | 2D CNN | 3D U-Net | 11 - 40M | 7 | Yes | MSE |
| 5 | N/A | 3D U-Net | 350K | 4 | No | CE |
| 6 | 2D CNN | 3D U-Net | 1.5 - 28 M | 10 | Yes | Focal Tversky |
| 7 | 2D / 3D CNN | 3D U-Net | 61 - 92 M | 6 | Yes | CE |
| 8 | N/A | 3D U-Net | 1.5 - 12 M | 24 | Yes | Dice, CE |
| 9 | 3D CNN | 3D U-Net | 12 - 24 M | 6 | No | CE |
| 10 | N/A | 3D U-Net | 13 M | 9 | Yes | Tversky, CE |

There were several innovations and adaptations beyond model architecture that the winners reported using (Extended Data Table 2). One common theme was the use of ensembles. Some winning teams implemented model soup^28^, an approach that averages the weights from independently-trained models with a shared architecture to improve prediction accuracy. Others, including the 1st place team, ensembled the intermediate predictions from models with distinct architectures, hypothesizing that increasing model diversity would yield superior results. A second common strategy incorporated extensive data augmentation during training, coupled with test time augmentation (TTA) during inference, to improve model generalization and prediction robustness (Extended Data Fig. 8). Many winning teams also increased the patch size used during inference compared to training, both to reduce border artifacts and to accelerate inference given the 12-hour runtime limit per submission. A fourth widely implemented strategy was model exponential moving average to make training more stable. Taken together, the winning designs were largely centered on overcoming the limited training data.

All winning teams reported ablation studies that quantified the contributions of features that yielded the final few percentage points in model performance (e.g. Table 3 in Uchida and Fukui^26^). These studies offered insights into which innovations catapulted these teams into the top 10 but did not explain the considerable gap in model performance between their solutions and particle picking methods developed in-field, including our benchmark solution. To investigate the origins of this gap, we started from a simplified version of our benchmark solution that consisted of a lightweight 3D U-Net with residual connections and minimal data augmentation performed during training. The resulting baseline achieved a model score of only 0.44. We then incorporated the innovations described by the winning teams into this minimal model to quantify how much each feature improved model performance. Increasing data augmentation during training alone had a dramatic impact on model performance, increasing the model score to 0.70 (Table 2). A non-negligible gap remains between this and the winning team’s score, which may require applying these strategies in combination or hyperparameter tuning to bridge. Nevertheless, these results highlight the importance of data augmentation under the realistic constraint of minimal training data.

**Table 2:** Sensitivity analysis quantifies the contribution of features used by the winning teams to model performance. The private test score of a minimal model consisting of a lightweight U-Net with residual connections was benchmarked as a baseline. Common features used by the winning teams were then incorporated into this baseline model, in a background that applied either minimal or expanded data augmentations during training (see Extended Methods).

| Model features | Augmentations |  |
| --- | --- | --- |
|  | Minimal | Expanded |
| Baseline | 0.44 | 0.70 |
| TTA (rotations) | 0.46 | 0.72 |
| TTA (rotations + flips) | 0.47 | 0.71 |
| Pretraining on synthetic data | 0.47 | 0.71 |
| Ensembles (model soup) | 0.56 | 0.73 |
| Increased inference patch size | 0.50 | 0.71 |
| Exponential moving average (EMA) | 0.43 | 0.69 |
| Training on beta-amylase | 0.55 | 0.72 |
| Larger model architecture | 0.38 | 0.69 |

### Additional highlights

This challenge yielded notable results beyond the scope of what was explicitly scored. One highlight was the high quality of the beta-amylase predictions made by the winning team. Although 60% of final submissions did not make predictions for this species, the 1st place team found that its inclusion during training improved the prediction accuracy for beta-galactosidase, which resembles beta-amylase from certain viewing angles. Despite spanning only 976 picks compared to the 2.5k in the ground truth labels, the winning team’s beta-amylase annotations yielded a higher resolution reconstruction than the ground truth map and featured better resolved secondary structure (Fig. 4a, Extended Data Fig. 3c). This result, along with the generally high quality of the winning teams’ annotations, has prompted us to consider releasing an updated version of the ground truth that cleans up the beta-amylase labels and incorporates valid particles for the other species found by the contestants but missing from the ground truth labels. The consensus picks from the winning teams offer a promising starting point for this revised ground truth (Extended Data Fig. 7a-b). However, in some cases these models were confused by the same non-target particles and contamination, so additional curation would still be needed (Extended Data Fig. 7c). An updated ground truth could also be extended to nucleosomes, proteasomes, and other species derived from the cellular lysate to further challenge annotation algorithms to generalize to diverse molecules (Extended Data Fig. 7c).

**Fig. 4:**
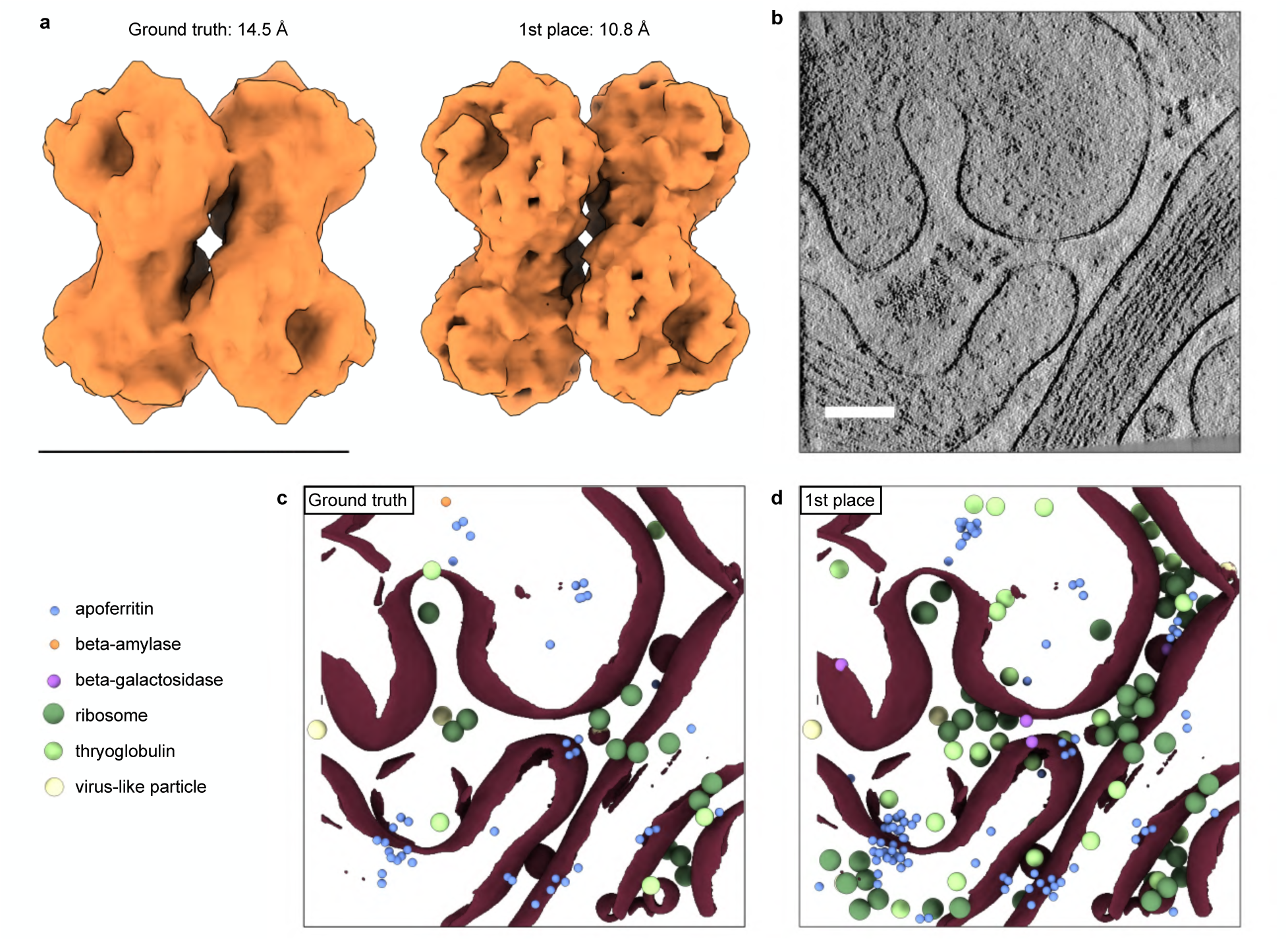
The winning team’s predictions outperformed the ground truth labels for beta-amylase and in crowded tomograms. **a.** Refined maps for beta-amylase from the ground truth and 1st place team’s annotations are compared. The scale bar corresponds to 100 Å. **b.** A slice from a private test set tomogram with significant molecular crowding is visualized. The scale bar corresponds to 100 nm. The **c.** ground truth labels and **d.** 1st place team’s annotations for the full volume are compared. Membrane segmentations^30^ are displayed in brown. The ground truth and 1st place team’s annotations are respectively available at CZCDP-10310 and CZCDP-10319 on the CryoET Data Portal^25^.

A second highlight is the robust performance of the top-ranked models against tomograms with significant molecular crowding. One limitation of the phantom dataset is that it does not emulate the extreme molecular crowding of cellular tomograms, a trade-off that made it feasible to rigorously annotate a large experimental dataset in a reasonable time frame. Though the training tomograms tended to be sparse, several test tomograms were characterized by crowding comparable to in situ data (Fig. 4b, Extended Data Fig. 9a). Examining the 1st place team’s annotations for these tomograms revealed that their model localized numerous valid particles that were missing from the ground truth labels even in regions full of cellular debris (Fig. 4c-d, Extended Data Fig. 9b-c). This observation suggests that these algorithms have the potential to generalize to cellular tomograms, a topic that will be critical to address in future work.

## Discussion

This competition successfully engaged >1000 experts from diverse domains to deliver new multi-class particle picking algorithms that outperformed existing state-of-the-art. Collectively the contestants’ results emphasized the importance of data augmentation and other strategies to overcome limited training data and underscored the detrimental impact of severe over-picking on STA. However, the relative insensitivity of STA to a moderate number of false positives exposes the difficulty of measuring pick accuracy, which in turn creates obstacles to comparing model performance and highlights the need for more robust evaluation metrics. Importantly, this competition also validated the phantom dataset^24^ as a useful benchmark for multi-class annotation algorithms. The design of this dataset — in particular, the limited training set and inclusion of small and easily confused targets — challenged algorithms to perform robustly under realistic constraints, while its considerable size of nearly 500 tomograms ensured rigorous evaluation against withheld data. Though Kaggle’s data formatting and time constraints made it prohibitive for many cryoET developers to participate, this dataset has been leveraged outside of the competition to benchmark new algorithms^29^ and refine existing tools^16^.

Two months after the challenge ended, we hosted a workshop that gathered members from the cryoET and Kaggle communities to discuss next steps. Despite widespread enthusiasm for a follow-up competition, consensus about its objective was difficult to reach. CryoET practitioners strongly favored a competition focused on unsupervised approaches to classify unlabeled densities in cellular data, with scoring based on STA. However, this runs counter to the Kagglers’ preference for competitions with delimited tasks and simple scoring metrics that enable rapidly iterating on algorithm design. With the full phantom dataset and winning submissions now released on the CryoET Data Portal^25^, we anticipate that the next several months will spur further innovations and offer insights into the capacity of the competition models to generalize to in situ data. These results may in turn highlight specific gaps in particle picking algorithms that the next challenge could be tailored to overcome.

## Extended Methods

### Description of the competition data

The phantom dataset was curated based on a combination of visual inspection and quality metrics as previously described^24^. This curation yielded 492 high-quality tomograms that were divided into training, public test, and private test sets. Dataset splits were designed to have similar distributions of particle ratios across the 6 annotation targets based on the Kolmogorov-Smirnov test. This analysis was used to group the dataset into 16 bins, which tomograms were randomly sampled from to construct the dataset splits. Experiments that tested training set sizes ranging from 2 to 87 tomograms indicated that a minimum of 7 tomograms was needed to match the distribution of the two test sets. Our best-effort annotation algorithm at the time, DeepFindET^24^, was trained on different subsets of 7 tomograms to confirm that this provided sufficient ground truth labels for successful model training. The 7 tomograms in the competition training set were manually selected from a set of 20 candidate tomograms. In addition to matching the test set distributions, these specific tomograms were chosen since they contained confusion factors such as membranes, contamination, strong edge artifacts, carbon film edge, and surface radiation damage/artifacts. The public test set comprised 25% of the remaining tomograms (121 total), which were withheld from the contestants. However, model scores for this split were made available on a public leaderboard that was continually updated on the competition’s webpage. The final 364 tomograms formed the private test set. Both the tomograms and model scores for this split were withheld since these private test scores were used to rank the final submissions.

For each tomogram in the training set, multiple versions were provided to the contestants as an optional source of data augmentation during training. These tomogram types, which shared alignments but differed in their SNRs, included: 1) weighted back-projection, which has the least contrast; 2) contrast transfer function (CTF)-deconvolved, which has intermediate contrast and was generated by applying AreTomo3’s local CTF correction to the tilt images prior to reconstruction; 3) denoised, which applied DenoisET to enhance the contrast of CTF-deconvolved tomograms; and 4) IsoNet-corrected, which applied IsoNet’s missing wedge restoration and denoising algorithm to the denoised tomograms^31,32^. Due to data storage constraints, both the public and private test sets consisted only of the denoised tomograms. For the competition, all training and test tomograms had a pixel size of 10 Å. After the competition ended, CTF-deconvolved and denoised tomograms at pixel sizes of 5 and 10 Å were released on the CryoET Data Portal for both test sets^25^.

To reduce barriers to entry, we provided example notebooks that demonstrated running DeepFindET^24^, TomoTwin^16^, and a generic 3D U-Net model on the competition data. These notebooks offered a workflow for training (except for TomoTwin, which relies on a pre-trained model); inference; and submitting solutions through Kaggle’s platform. In addition, these notebooks illustrated using PyTorch-based functions from the copick library^33^ to handle cryoET metadata and data. These notebooks, the leaderboards, the winners’ reports, and other material are available on the competition’s website: https://www.kaggle.com/competitions/czii-cryo-et-object-identification. Links to the winners’ reports and deposition IDs for their annotations are also listed in Extended Data Table 2.

### Evaluation metric to score submissions

The ground truth annotations were meticulously curated by a team of cryoET experts but are necessarily imperfect since the phantom dataset is an experimental benchmark^24^. Though rigorous curation minimized the number of false positives, it was impossible to guarantee that valid particles were not missed. To account for this, contestants’ annotations for each particle class were evaluated based on the F_β_ score:

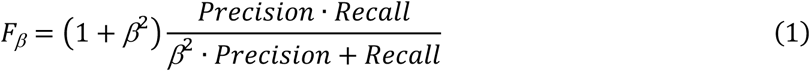

where the parameter, β, permits tuning the relative extent to which precision:

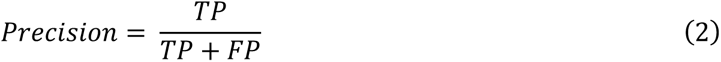

and recall:

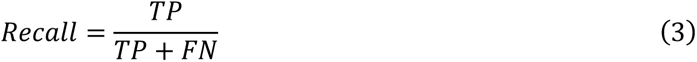

are penalized. TP, FP, and FN correspond to true positives, false positives, and false negatives, respectively. Submissions were scored with β=4 to apply a higher penalty to contestants’ FN than their FP in case the latter contained valid particles missing from the ground truth annotations. The submitted annotations were classified as TP if their coordinates were within a distance threshold of a ground truth annotation. The distance thresholds were set to half of each particle’s radius, resulting in thresholds of 30 Å for apoferritin, 32.5 Å for beta-amylase, 45 Å for beta-galactosidase, 75 Å for ribosomes, 65 Å for thyroglobulin, and 67.5 Å for virus-like particles. If multiple submitted coordinates were within the threshold distance of a ground truth annotation, only one was scored as a TP and the remainder were considered FP.

Submissions were assigned an aggregate score based on the weighted average of the per-particle F_β_ scores:

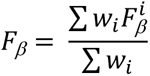

where the virus-like particle, ribosome, and apoferritin classes were assigned a weight (*w*) of 1, while beta-galactosidase and thyroglobulin were assigned a weight of 2. Though ground truth labels for beta-amylase were provided for the training data, this particle class was not scored due to insufficient confidence in our annotation accuracy for this species. As noted above, model scores for the public test set were made available throughout the competition, while the private test scores were only released after the competition ended. Extended Data Fig. 1b shows the correlation between the public and private test scores across all valid submissions.

### Comparison to approaches developed in-field

DeepFinder^13^ and DeepETPicker^17^ were benchmarked against the competition data to compare the contestants’ performance to algorithms developed and commonly used by cryoET practitioners. For both software packages, 7 independent models were trained on the training set, with a different training tomogram held out for cross-validation during each run. The ground truth beta-amylase labels were not used during training. DeepFinder offers the option to model annotation targets using spherical or shape-based masks; for this analysis, the sphere-based model was chosen since the shape-based approach requires orientation information, which was not provided to the contestants. For both algorithms, label sizes for the ground truth annotations were based on each species’ dimensions, and the default hyperparameters were used during training and inference. Once trained, inference was performed by applying each model separately to the public and private test sets. The variation in private test scores across the 7 runs was small for DeepETPicker (0.68±0.01). In the case of DeepFinder, only 3 of the 7 runs yielded particle predictions and had a mean private test score of 0.54±0.02. To mimic the contestants’ use of the public leaderboard scores for model selection, the private test score of the model that yielded the best public test score is reported in Fig. 1d, and the annotations predicted by this selected model were used for STA. While hyperparameter tuning would likely improve these algorithms’ performance on the phantom dataset, such experiments were beyond the scope of this work. Consequently, we limited our analysis to how these in-field methods performed out-of-the-box but acknowledge that the comparison favors the Kaggle contestants, many of whom performed extensive hyperparameter tuning during the three months of the competition.

### Generation of consensus and removal of ground truth

The consensus picks were generated from the common annotations among the top 5 teams. For each species, coordinates between teams were considered matches if their inter-particle distance was less than the species’ radius, and the coordinates for matching picks were averaged. Consensus picks could not be generated for beta-amylase since only one of the top 5 teams submitted predictions for this species. To assess the quality of the contestants’ FP and estimate the extent to which the ground truth coordinates were under-picked, the individual team’s and consensus annotations were reanalyzed after removing the ground truth labels from each set of picks. This was achieved by omitting coordinates that matched a ground truth annotation using each species’ radius as the distance threshold. 2D analysis of the consensus picks after ground truth removal was performed by generating per-particle projections using the slabpick library, followed by classification of these particle projections in CryoSPARC^34^ as previously described^24^.

### Subtomogram averaging

The ground truth and contestants’ coordinates were stored in copick^33^ format, which streamlines parallel access of cryoET annotations and tomograms. These coordinates were converted to RELION-5^27^ particle STAR files using py2rely, a software package that integrates the CCPEM-pipeliner’s job execution functions (https://ccpem-pipeliner.readthedocs.io/en/latest/index.html) into a unified workflow to enable fully automated STA in RELION-5. The workflow followed the same approach applied to the ground truth labels as previously described^24^ with the following two exceptions. First, all STA results presented in this work are from the annotations for the combined public and private test sets and exclude coordinates for the training set. Second, neither Bayesian polishing nor CTF refinement was performed. For each particle class, the same reference was used for STA across all teams, and no particles were discarded -- for example, through classification or any other type of curation -- to ensure consistency when comparing the teams’ picks. Duplicate particles were also retained so that the refined maps reflected the full set of picks from each team.

Each set of annotations was processed using py2rely’s reference-based STA pipeline, which automated the following steps. First, the picks from each tomogram were evenly and randomly split between half sets. Next, initial angular orientations were assigned by running a 3D classification job at bin 6 (9.06 Å/pixel) with a single class against templates from the following reference structures: EMD-41923 (apoferritin), EMD-0153 (beta-galactosidase), EMD-30405 (beta-amylase), EMD-3883 (ribosome), EMD-24181 (thyroglobulin), and EMD-41917 (virus-like particle). Templates were low-pass filtered to 20 Å for apoferritin and to 40 Å for all other species. Particle orientations were refined at the bin 6 resolution and then re-extracted at bin 2 (6.02 Å/pixel). A second round of refinement was performed using the reconstructed map as the reference and a particle-shaped mask, which was low-pass filtered to 10 Å and padded by 3 pixels with a 5 pixel-wide soft-edge. For the apoferritin, ribosome, and virus-like particle species, particles were re-extracted at bin 1 (1.51 Å/pixel) and a final refinement was performed. Bin 1 re-extraction and refinement were skipped for the other species, which did not achieve Nyquist resolution at bin 2. In the case of apoferritin, processing -- including 3d classification with a single class and refinement -- started at bin 2 before proceeding to the bin 1 steps described above.

Map resolution was calculated as the spatial frequency at which the map-model Fourier shell correlation (FSC) decreased to 0.5^35^. The map-model rather than half-maps FSC^36^ was computed to avoid the latter’s sensitivity to symmetrization artifacts (Supplemental Note 2). Reference maps were generated using ChimeraX’s^37^ *molmap* function from the following atomic models from the Protein Data Bank^38^: 1FA2 (beta-amylase), 6DRV (beta-galactosidase), 6QZP (ribosome), and 7B75 (thyroglobulin). For apoferritin, the reference map was derived from an Electron Microscopy DataBank^39^ map (EMD-19880). For the virus-like particle, a reference map from a single-particle analysis study was provided by collaborators at NYSBC, who also supplied the stock virus-like particles used in the phantom sample. In these two cases, an experimental rather than simulated cryoEM map was used to improve modeling of the solvent interior of these shell-like proteins, which the *molmap* function treats as vacuum (Supplemental Note 2).

For each particle class, the same mask was applied for all FSC calculations to ensure an unbiased comparison between sets of annotations. All masks were derived from density simulated from fitted atomic models using ChimeraX’s *molmap* function rather than the mask used during refinement, since the latter was generated as part of refinement and consequently was not consistent between different sets of picks. For beta-amylase, beta-galactosidase, and thyroglobulin, a mask that covered the full protein was applied. For the ribosome, a mask was applied so that the FSC calculation was based exclusively on the large subunit to reduce the impact of inter-subunit ratcheting on the resolution estimate. For apoferritin and the virus-like particles, a mask was applied to isolate the comparison to a single asymmetric unit. Independent STA runs were carried out in triplicate for four of the species (apoferritin, beta-amylase, thyroglobulin, and virus-like particles) to quantify the variability in map quality due to nondeterministic behavior of RELION-5’s STA pipeline (Supplemental Note 2). In these cases, resolution estimates are reported for the run that yielded the highest resolution map. For each species, maps are displayed with the isosurface threshold set to the following multiple of the standard deviation from the map’s mean value unless otherwise noted: apo-ferritin: 4, beta-amylase: 5, beta-galactosidase: 5, ribosome: 3.5, thyroglobulin: 4, and virus-like-particle: 3.5.

### Sensitivity analysis

The relative contributions of different features to model performance were evaluated using Octopi, a modular deep learning framework for particle picking (see Code Availability). Tests were performed using the same model architecture that was used for the benchmark solution shown in Fig. 1b, which consisted of a U-Net architecture with a ResNet backbone composed of four encoding layers and one residual block per layer. Considering this model architecture as the baseline, we assessed the following features individually and in combination:

- Additional data augmentation, which included intensity adjustments, random cropping, the addition of Gaussian noise, rotations of 90° in the xy plane, and 180° flips along each axis. By contrast, the default data augmentations did not include flips, performed Gaussian smoothing rather than adding Gaussian noise, and applied intensity adjustments with a higher probability, resulting in few training examples with unaltered contrast compared to the original data.
- Test-time augmentation (TTA), which applied 90° rotations in the xy-plane with uniform probability during inference and then made class assignments by averaging predictions from the augmented data. A separate experiment applied these rotations in combination with 180° flips along the x and/or y axes.
- Pretraining on synthetic data, after which the model was fine-tuned on the training set. The synthetic data, which contained the six annotation targets and membrane structures, were simulated using PolNet^40^ and provided as an optional resource to contestants during the competition.
- Ensembling via model soup^28^, in which the model weights from the 3 top-performing models of 7 independently-trained models were averaged.
- Increased patch size during inference, with the patch side length doubled to 160 voxels during inference compared to the default value.
- Exponential moving average (EMA) of the model weights, with a decay rate of 0.99.
- Training with beta-amylase, in which the ground truth labels for beta-amylase were included during training.
- A larger model architecture, which contained one more encoding layer and one additional residual block per layer compared to the baseline model. This larger model contained 2.7x more parameters than the baseline model’s 1.1 million parameters.

For each experiment, the model was trained for a maximum of 3000 epochs using the Tversky Loss (α=0.3, β=0.7) and Adam optimizer with one training tomogram held out for cross-validation. The epoch that achieved the maximum F_β_ score on the withheld tomogram was selected for inference. During inference, the predicted segmentation masks were converted to picks using the Watershed algorithm to separate clusters, followed by a size-based selection scheme to convert clusters that ranged in size from half to the full particle’s radius to individual coordinates. Three independent replicates with 7-fold cross-validation were run per experiment to assess the variation in model performance due to stochastic weights initialization and random sampling from the training data. On average, 12% of runs failed to predict coordinates for at least one target species. Omitting these failed runs reduced the standard deviation in the private test scores between replicates for each experiment from 0.05 to 0.01, indicating minimal variation due to nondeterministic behavior. As with DeepFinder and DeepETPicker, the private test scores reported in Table 2 are based on the model that achieved the highest public test score for the indicated trial to mirror the contestants’ use of the public leaderboard scores for model selection. Our best-effort model shown in Fig. 1b is the baseline model in combination with pre-training on synthetic data, including beta-amylase labels during training, and increasing the patch size used during inference compared to training.

## Data Availability

The phantom dataset is available on the CryoET Data Portal under deposition ID CZCDP-10310 (https://cryoetdataportal.czscience.com/depositions/10310) and divided among the training set (DS-10440), the public test set (DS-10445), and the private test set (DS-10446) according to the splits used during the competition. This deposition also contains the ground truth labels, annotations from the 10 winning teams, and a synthetic dataset (DS-10441) designed to mimic the phantom data. Maps from the ground truth labels and the 1st place team’s annotations have been deposited on the EMDB. For the ground truth, the deposition IDs are apoferritin: 73634, beta-amylase: 73640, beta-galactosidase: 73638, ribosome: 73631, thyroglobulin: 73642, and virus-like particle: 73636. For the 1st place team, the deposition IDs are apoferritin: 73635, beta-amylase: 73641, beta-galactosidase: 73639, ribosome: 73633, thyroglobulin: 73643, and virus-like particle: 73637.

## Code Availability

Octopi is available at https://github.com/chanzuckerberg/octopi. The version of py2rely used in this work is available at https://github.com/chanzuckerberg/py2rely/tree/kaggle.

## Acknowledgements

We thank the Kagglers who participated in our competition and especially the top 10 teams for their presentations and competition reports. We also thank M. Kopylov from NYSBC for providing an experimental reference map of the virus-like particle. We acknowledge the following individuals from the Chan Zuckerberg Initiative (CZI) SciTech team for contributing to the CryoET Data Portal, which supported the competition data and related material: A. Sweet, B. Nelson, E. M. Wang, R. Agarwal, T. Smith, B. Chu, D. Sadgat, E. Hoops and J. Larsen. K. Maitland and S. Otte from CZI supported the planning and execution of the competition. CZ Imaging Institute is made possible with support from Chan Zuckerberg Initiative (CZII-2023–327779).

## Author contributions

M.P. and K.H. designed and directed the Kaggle challenge. W.R. led hosting the challenge on Kaggle. J.S., S.K., Z.Z., and K.H. provided example notebooks for the competition. J.A., K.E., J.G., K.K., S.M., D.G.M., J.X.N., J.P., M.V., and Z.S.W. generated the competition webpage and other materials. B.A., D.L., and S.A.S.J. participated in the challenge and provided perspective on contestants’ models. Y.Y., A.P., and M.P. analyzed intermediate results from the challenge. J.S. U.H.E., D.K., S.Z., and Z.Z. analyzed the contestants’ models. J.S. and J.H. developed the STA pipeline. J.H. and A.P. performed the structural analysis. J.S. and A.P. performed the sensitivity analysis. A.P. wrote the manuscript with advice from M.P. and J.H. All authors reviewed the manuscript. D.A.A., C.S.P, and B.C. provided overall leadership for this work.

---

**Extended Data Fig. 1:**
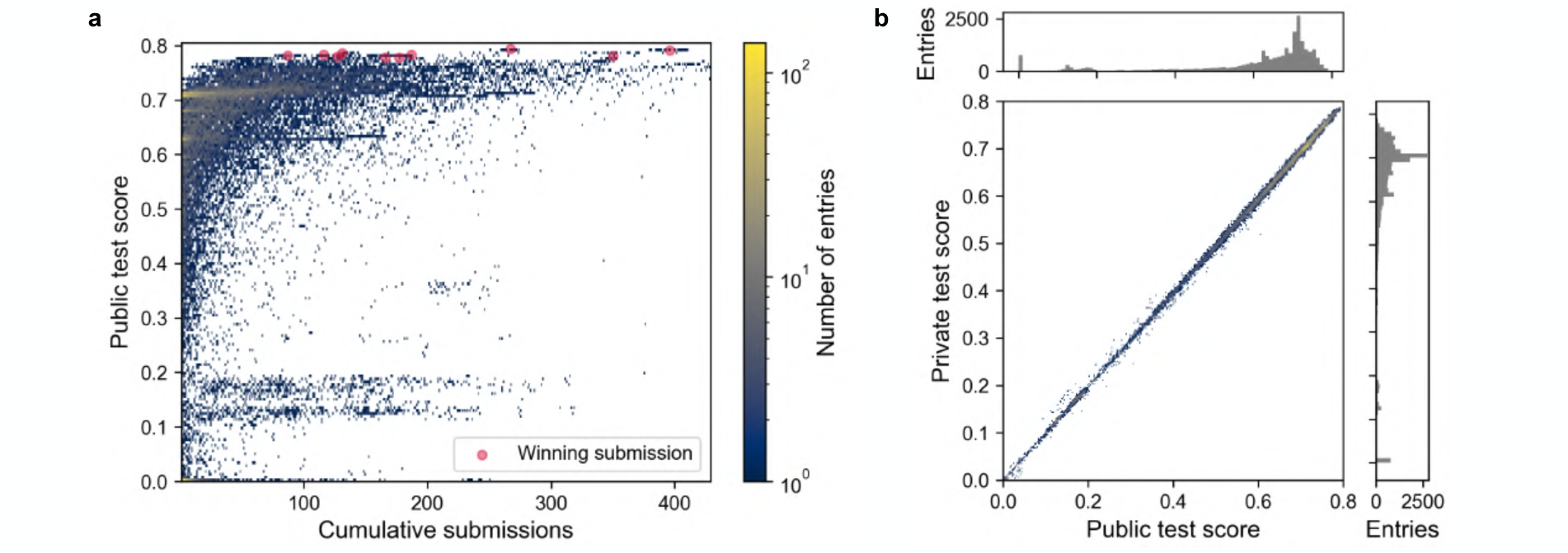
The evolution of public test scores and their correlation with private test scores. **a.** The cumulative number of entries submitted by each team is plotted as a function of the submission’s public test score. The red circles mark the final submissions from the 10 winning teams. **b.** The high correlation between the public and private test scores across all submissions indicates that the tomograms were successfully divided into subsets with similar particle distributions.

**Extended Data Fig. 2:**
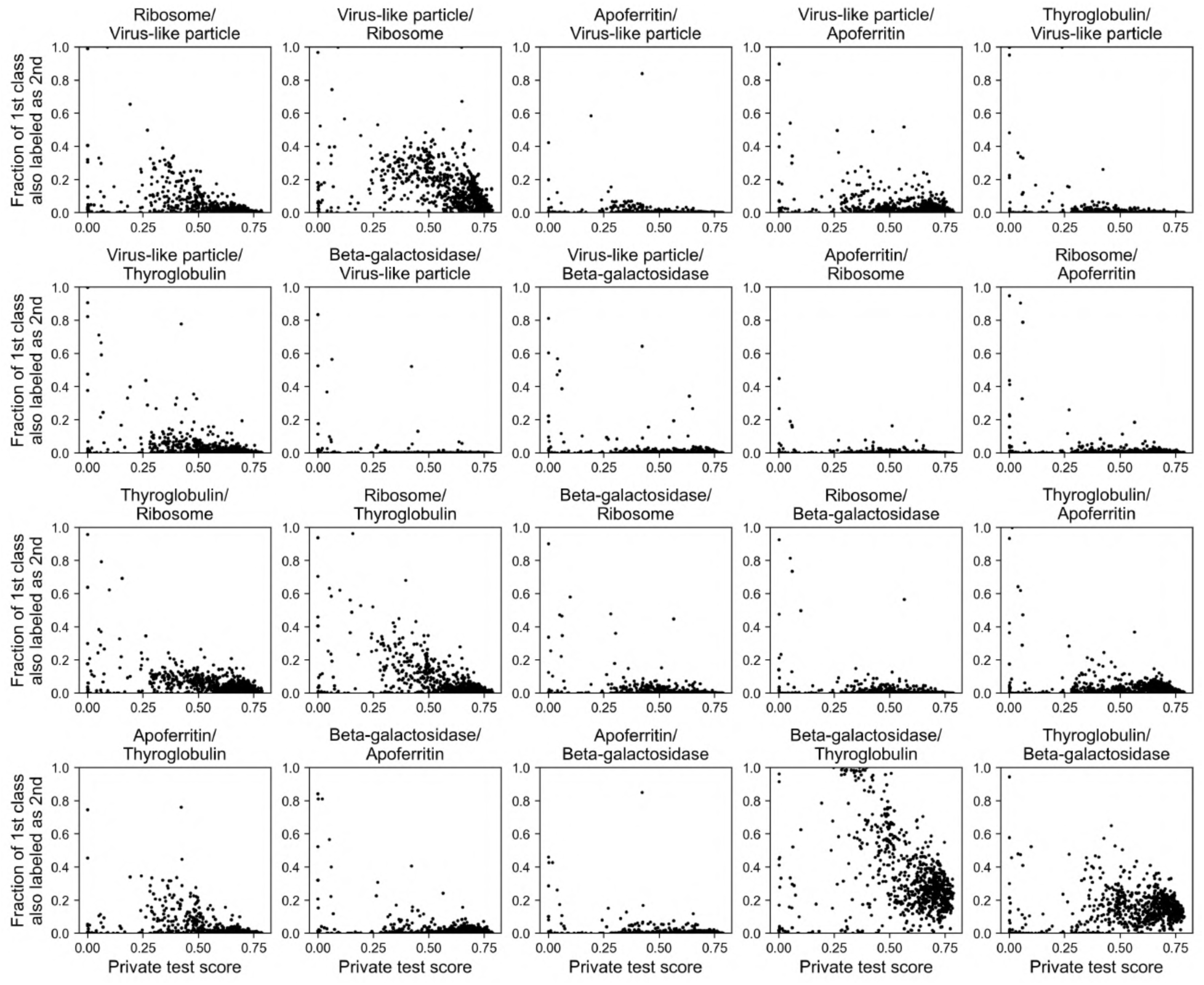
Analysis of inter-class duplicates indicates that higher-scoring models better discriminated between species. Each panel plots the fraction of coordinates assigned to multiple particle classes as a function of the submission’s private test score. Particle coordinates in the first-listed class were considered a match or duplicate if any coordinates in the second particle class were located within the first particle’s radius.

**Extended Data Table 1:**
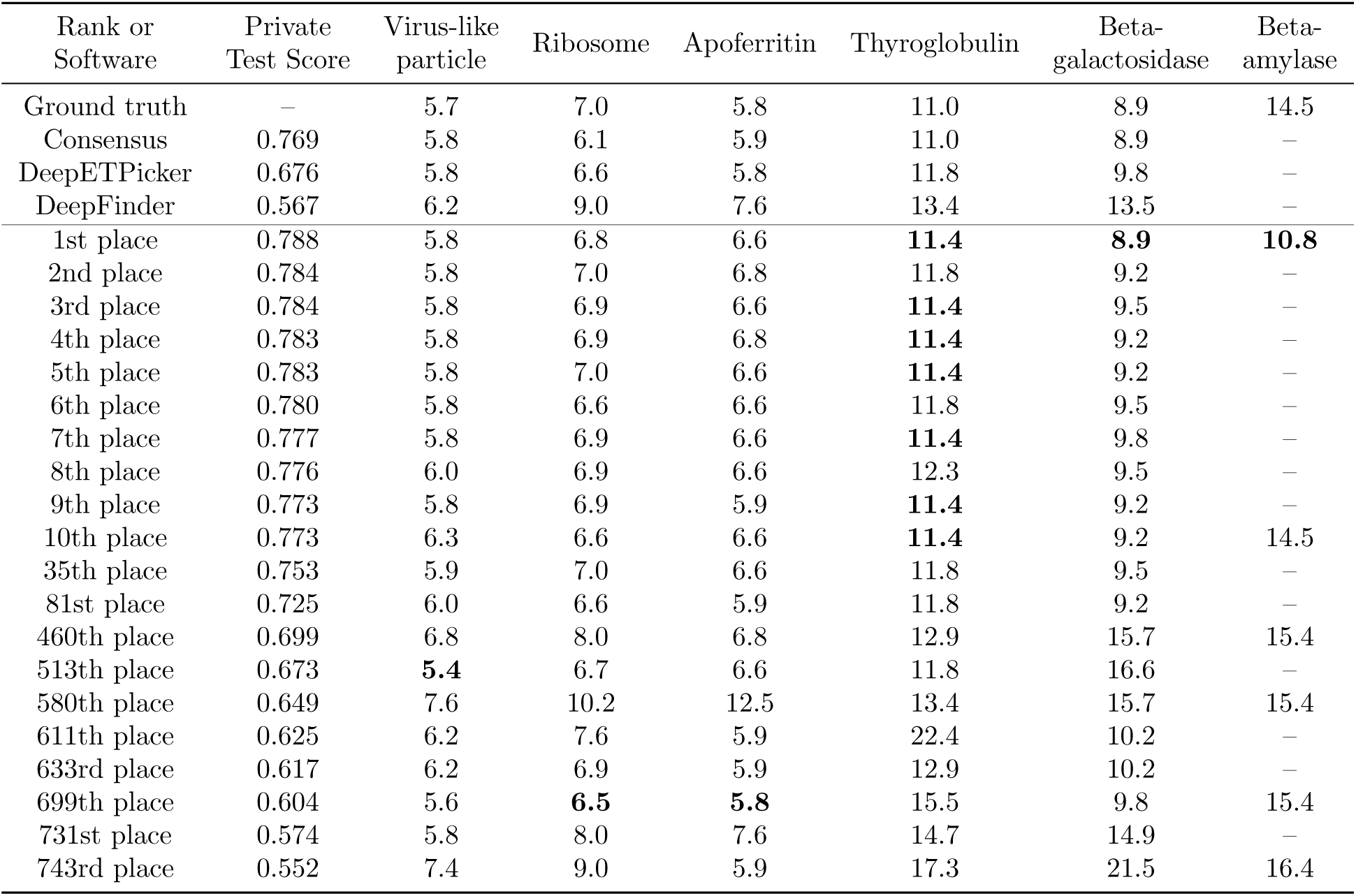
Comparison of STA results. Resolution estimates are based on the map-model FSC and reported in Å. Most of the teams assessed did not submit predictions for beta-amylase. Bold entries correspond to the highest resolution map from the team submissions for each particle class. For the virus-like particle, apoferritin, thyroglobulin, and beta-amylase classes, the resolution is reported for the run from three independent STA trials that yielded the highest resolution map. For intermediate-to high-quality maps (within 3 Å of the ground truth map), the standard deviation between replicates was on average <0.2 Å for the virus-like particle, apoferritin, and thyroglobulin classes. Inter-run variability was generally larger for poor quality maps as described in Supplemental Note 2. The standard deviation of the 1st place team’s beta-amylase replicates was 0.8 Å.

**Extended Data Fig. 3:**
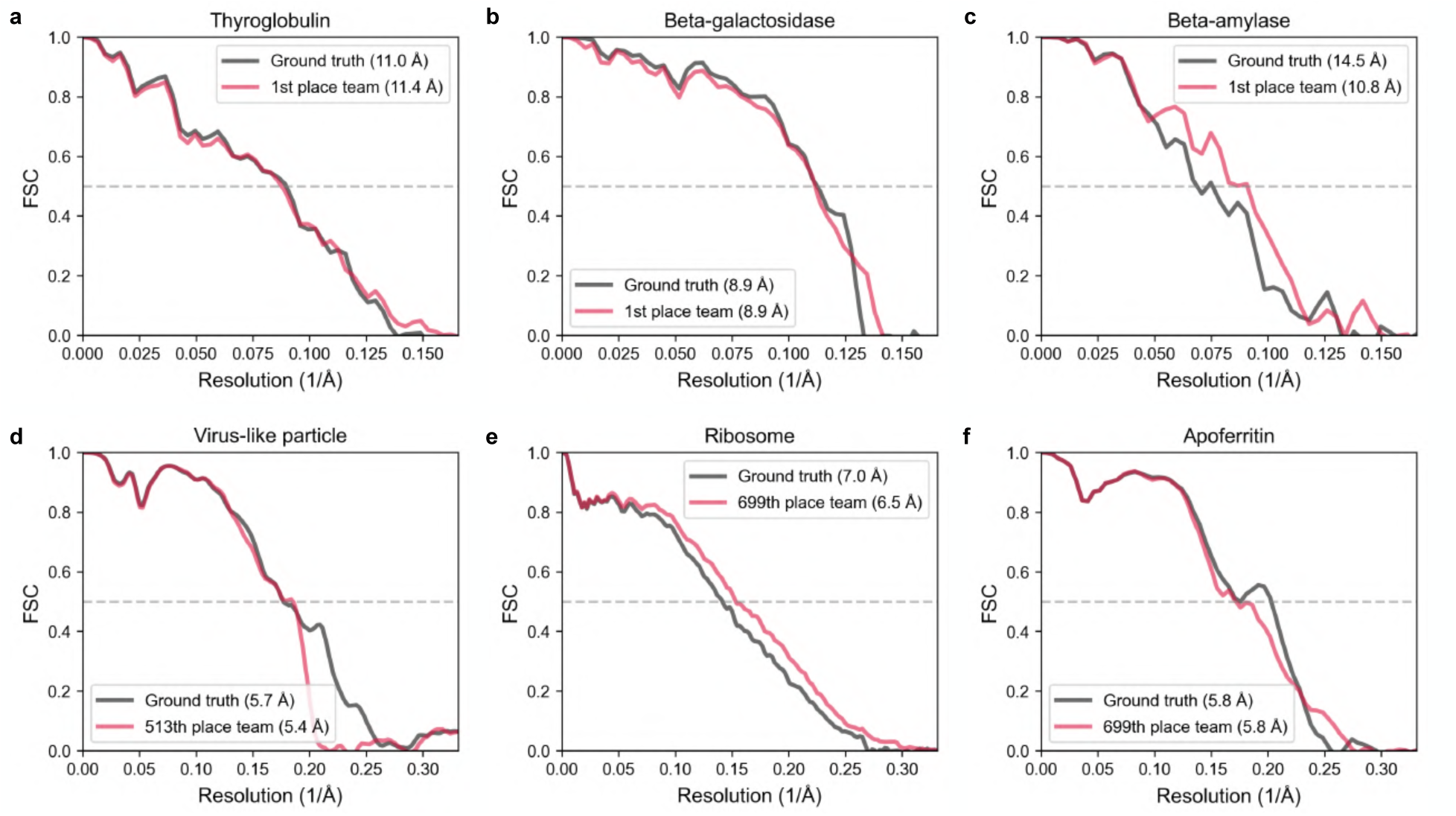
FSC comparison between the ground truth map and the highest resolution map from the Kaggle teams for each annotation target. The map-model FSC curves are compared for the ground truth and the Kaggle team that obtained the highest resolution map for **a.** thyroglobulin, **b.** beta-galactosidase, **c.** beta-amylase, **d.** virus-like particle, **e.** ribosome, and **f.** apoferritin. The 1st place team achieved the highest resolution map for the difficult targets, though in the case of thyroglobulin, multiple teams in the top 10 (but none of the lower-ranked teams) also obtained 11.4 Å maps. For the easy annotation targets, lower-ranked teams produced the highest resolution map.

**Extended Data Fig. 4:**
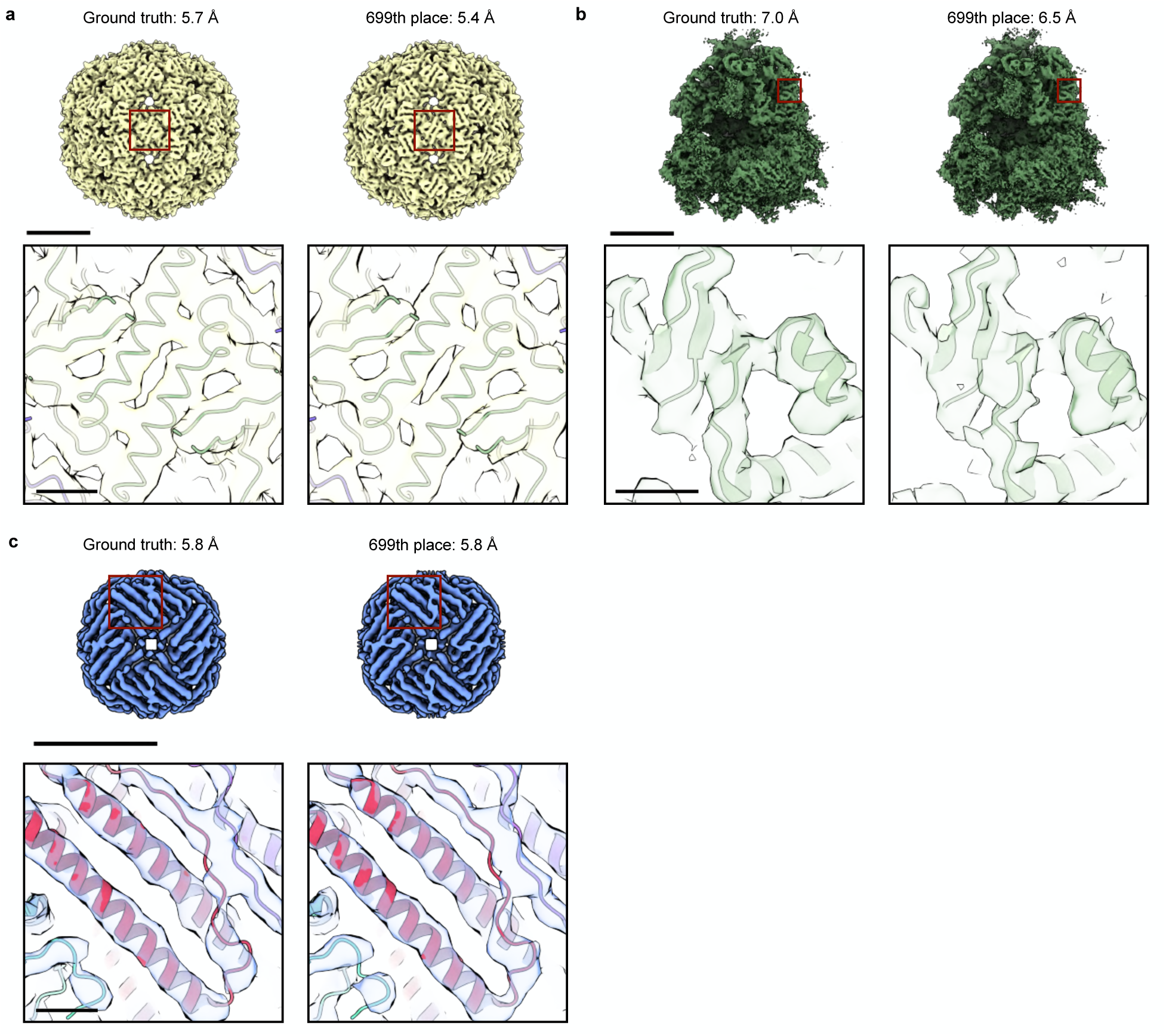
Structural comparison between the ground truth map and the highest resolution map from the Kaggle teams for the easy annotation targets. Maps from the ground truth labels and the annotations from the team that achieved the highest resolution are compared for the **a.** virus-like particle, **b.** ribosome, and **c.** apoferritin classes. For each species, the upper panel displays the full structure. The scale bar indicates 100 Å. The regions boxed in red are visualized in the insets below and overlaid on a reference atomic model. The scale bar indicates 10 Å. In all three cases, annotations from a Kaggle team ranked outside the top 10 yielded the highest resolution map.

**Extended Data Fig. 5:**
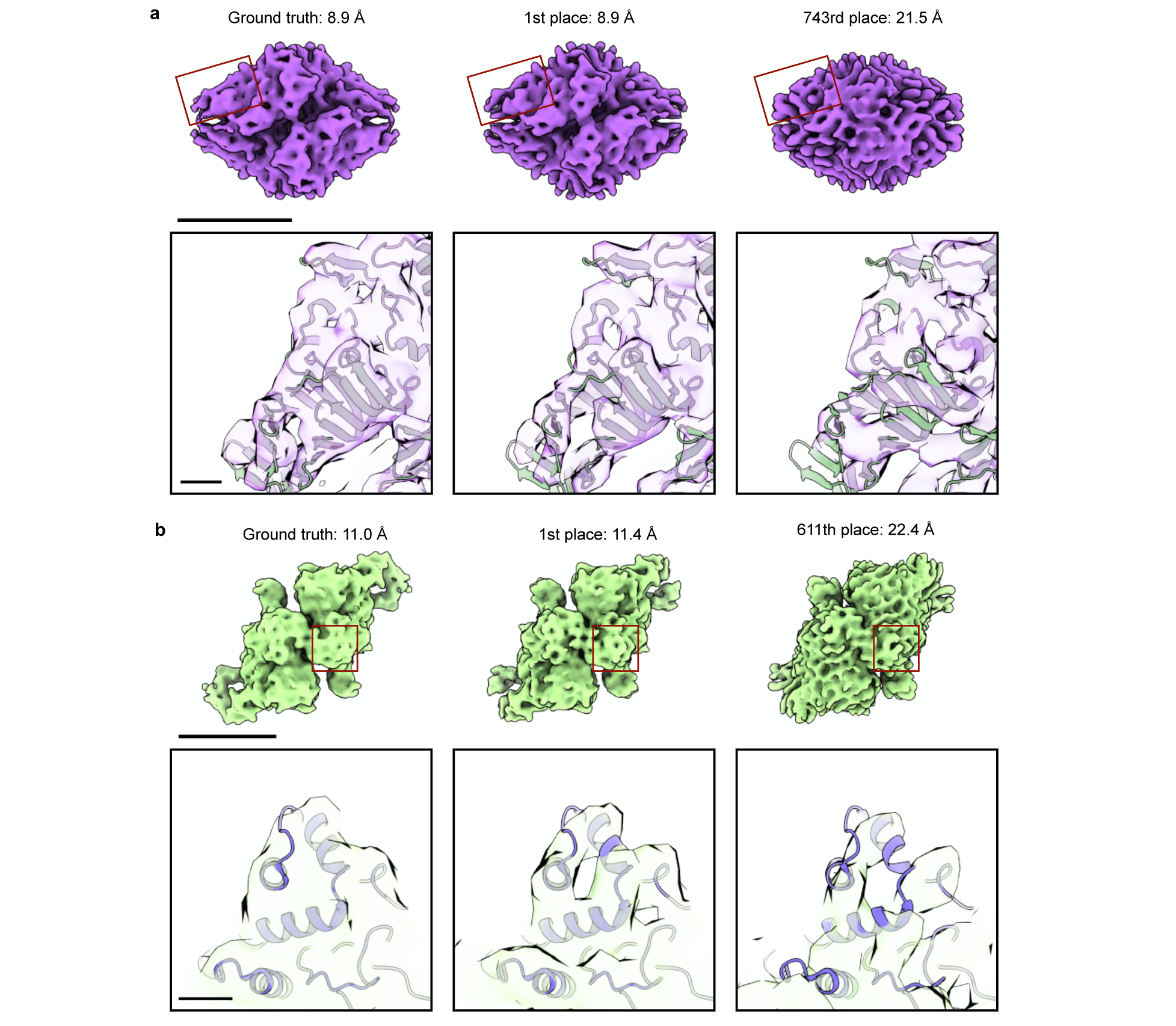
Structural comparison between the ground truth and the highest and lowest resolution maps from the Kaggle teams for the difficult annotation targets. Maps from the ground truth labels (left column) and the teams that achieved the highest (center column) and lowest (right column) resolution are compared for **a.** beta-galactosidase and **b.** thyroglobulin. The upper panel displays the full structure. The scale bar indicates 100 Å. The regions boxed in red are visualized in the insets below and overlaid on a reference atomic model. The scale bar indicates 10 Å. In the case of beta-galactosidase, the highest resolution map from the Kaggle teams was achieved by the 1st place team. In the case of thyroglobulin, annotations from several top 10 teams but none of the lower-ranked teams achieved 11.4 Å maps. Symmetrization artifacts are apparent in the lowest resolution maps for both species (right column).

**Extended Data Fig. 6:**
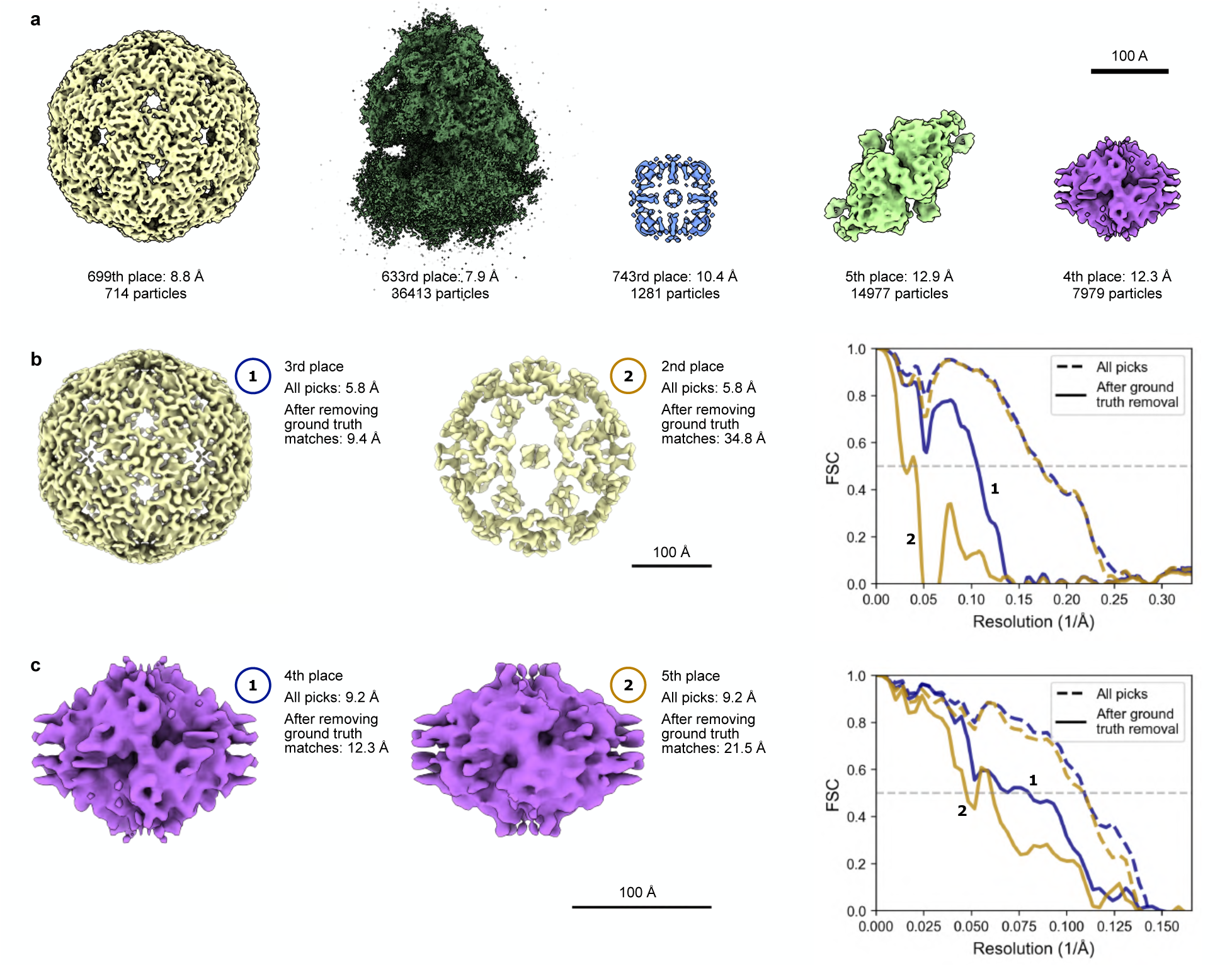
Comparison of map quality from the teams’ full set of annotations and their presumed false positives. To assess the quality of each team’s false positives, annotations that matched the ground truth labels were removed and the remaining picks were run through an automated STA pipeline. **a.** The highest resolution map from the teams’ presumed false positives is visualized for each species. With the exception of apoferritin, multiple teams were able to recover the correct structure even after removing the true positives. In the case of apoferritin, all other teams’ maps than the one shown were lower resolution despite having an order of magnitude more particles on average, indicating that particle counts were not resolution-limiting. **b-c.** In multiple cases, picks from teams that achieved the same resolution yielded maps of significantly different quality after the ground truth matches were removed from the annotation set. (left, center) Maps after ground truth removal are compared. The 34.8 Å virus-like particle map was characterized by abnormal intensity statistics, so the isosurface threshold was set to 95% of the maximum pixel value for visualization. (right) FSC curves are plotted for the indicated maps from the full annotation set (dashed lines) and from the presumed false positives (solid lines).

**Extended Data Fig. 7:**
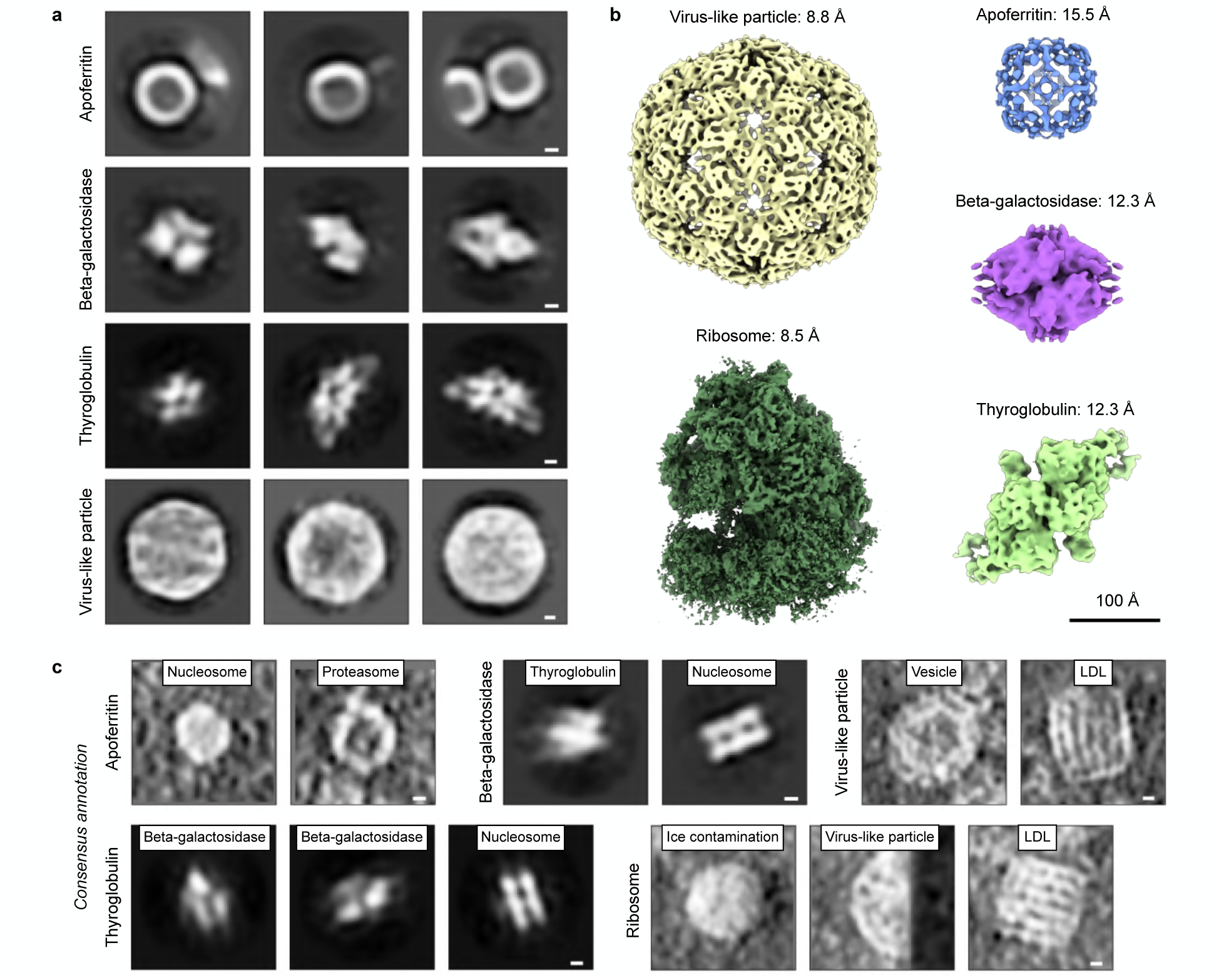
After omitting particles that matched the ground truth labels, the remaining consensus picks contained both valid particles and identifiable false positives. **a.** Valid particles present in the consensus picks from the top 5 teams but missing from the ground truth annotations were observed in 2D class averages of per-particle projections from each species (see Methods). The numbers of particles that sorted into correct classes (as judged by visual inspection) were apoferritin: 11,099, beta-galactosidase: 2624, thyroglobulin: 9931, and virus-like particle: 238. This provides a rough estimate of the extent to which the ground truth was under-picked. The poor centering of the apoferritin particles may account for the low quality of this species’ refined map. 2D class averages for the ribosome contained artifactual streaks, but the quality of its refined 3D map indicates that valid particles were present among the picks. The scale bar corresponds to 20 Å. **b.** Refined maps from the consensus picks after ground truth removal show lower resolution but recognizable maps for all species except for apoferritin. Compared to the ground truth map, resolution worsened by virus-like particle: 3.1 Å, ribosome: 1.5 Å, apoferritin: 9.7 Å, beta-galactosidase: 3.4 Å, and thyroglobulin: 1.3 Å. **c.** Identifiable contamination was also found among the 2D class averages. The consensus annotation is indicated on the left, while a putative label is noted at the top of each class. LDL is low-density lipoprotein. Scale bars correspond to 20 Å.

**Extended Data Table 2:**
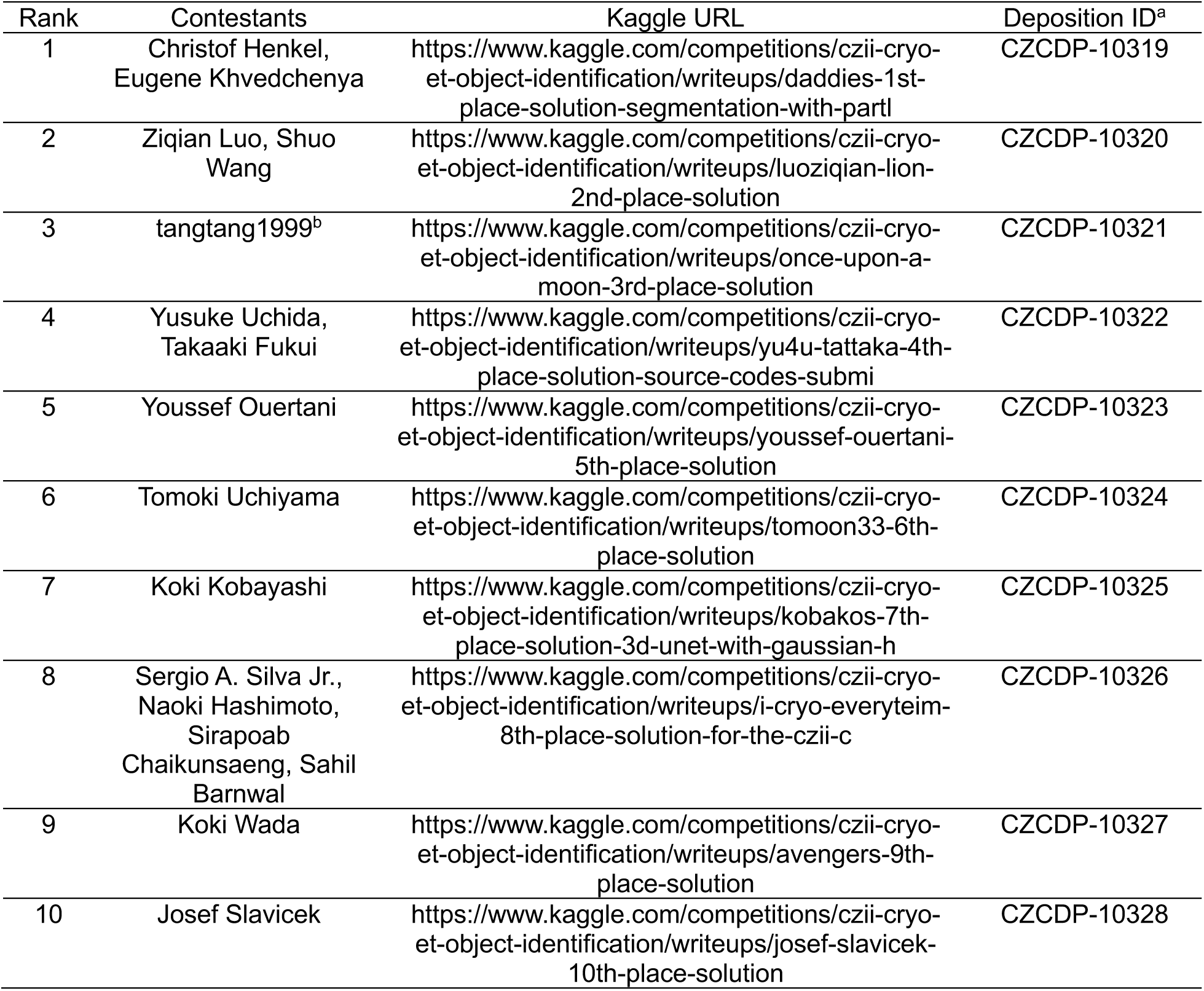
The winners’ published reports and annotations. The 10 winning teams released reports that described their models and training and inference procedures in depth. These are all available on the competition page hosted by Kaggle. ^a^The associated annotations have been published on the CryoET Data Portal^25^ under the listed deposition IDs. ^b^This contestant preferred for their annotations to be published under their Kaggle user name.

**Extended Data Fig. 8:**
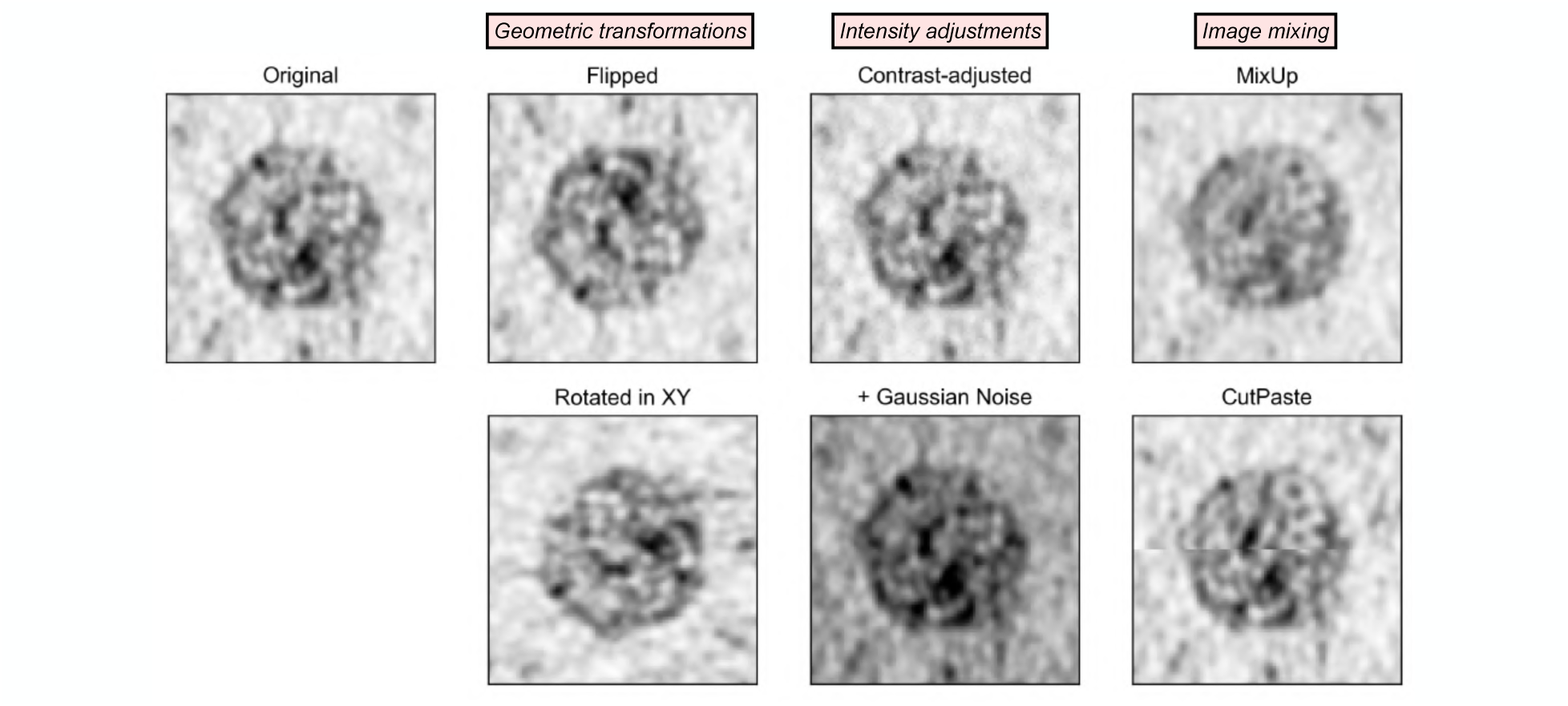
Types of data augmentation frequently used by the winning teams. All winning teams reported applying geometric transformations, which generally included both flips along the three tomogram axes and 90° rotations restricted to the XY plane (due to the missing wedge). Many also applied intensity adjustments, such as contrast adjustment or the addition of Gaussian noise. Some contestants used less conventional augmentations that mix images and their corresponding labels, including MixUp and CutPaste. While MixUp generates augmented training examples from the weighted average of subvolumes extracted from the tomograms, CutPaste creates new training data by cutting part of one training subvolume and pasting it into another. The latter augmentation was found to work well within but not between training tomograms since the tomograms’ different contrast levels led to more noticeable edge artifacts after pasting. Both types of image mixing augmentations were generally performed on training examples from the same particle class. In this figure, these augmentations were applied to a virus-like particle from the training data, and the resulting 2D projections are visualized. For reference, the original training example is displayed on the left. For the image mixing augmentations, the augmented examples were generated by combining two virus-like particle subvolumes from the same tomogram in a 50:50 blend.

**Extended Data Fig. 9:**
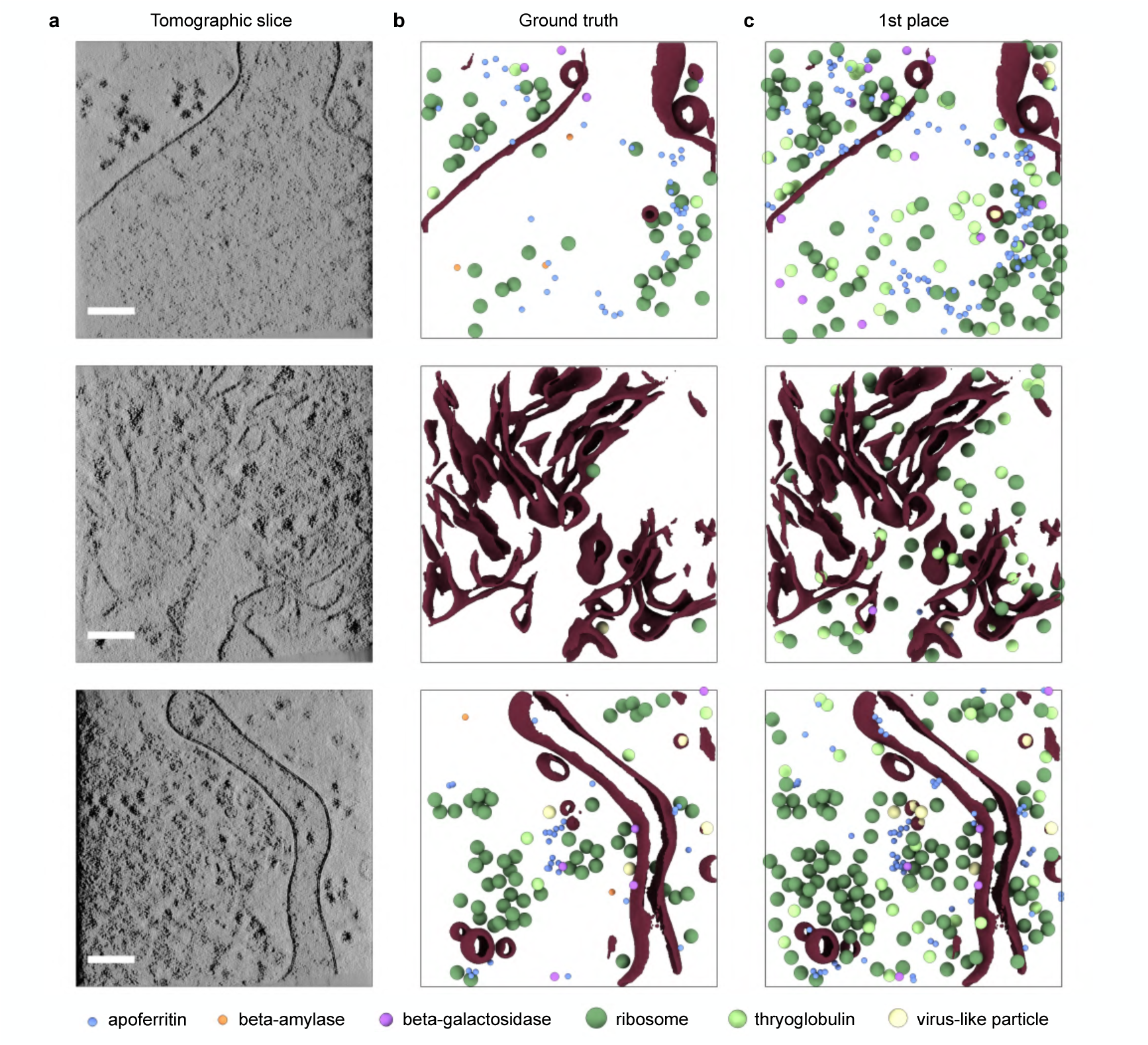
The winning team’s annotations were robust and more complete than the ground truth labels in crowded tomograms. **a.** Slices through private test set tomograms that feature significant molecular crowding are visualized. The scale bar indicates 100 nm. The **b.** ground truth labels and **c.** 1st place team’s annotations are shown for the full volumes. Membrane segmentations are displayed in brown.

## Notes

https://cryoetdataportal.czscience.com/depositions/10310

### Competing Interest Statement

The authors have declared no competing interest.

### Summary of Updates

Extended Data Table 2 has been added to cite the winning teams' reports and annotations, which are published on Kaggle's competition webpage and the CryoET Data Portal, respectively.

