## Supplemental Note 1 for "Lessons learned from a Kaggle challenge for particle picking in cryo-electron tomography"

### Supplemental Note 1: Comparison of refined maps for the challenging targets

Supplemental Figs. 1-3 compare the refined maps for the challenging annotation targets. From left to right, the top row visualizes the reference map generated from an atomic model with the density simulated to 6 Å resolution, the ground truth map, a map from the consensus picks from the top 5 teams, and maps from the DeepETPicker[17] and DeepFinder[13] annotations. The second and third rows display maps from the top 10 teams, and the remaining rows show maps from 10 lower-ranked teams. The listed score is the private test score for each set of annotations.

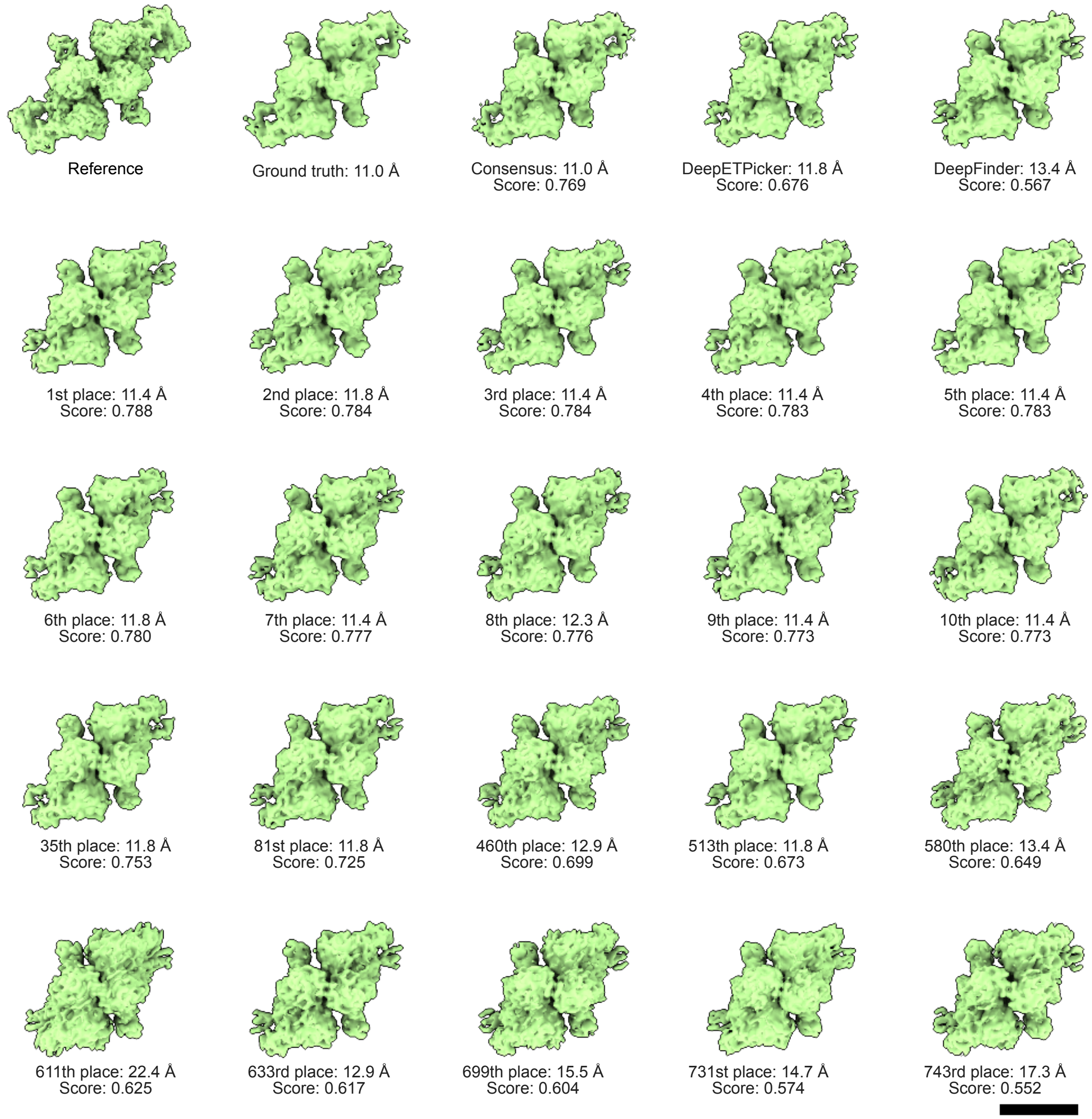


**Supplemental Fig. 1: Comparison of thyroglobulin maps**. The scale bar corresponds to 100 Å.

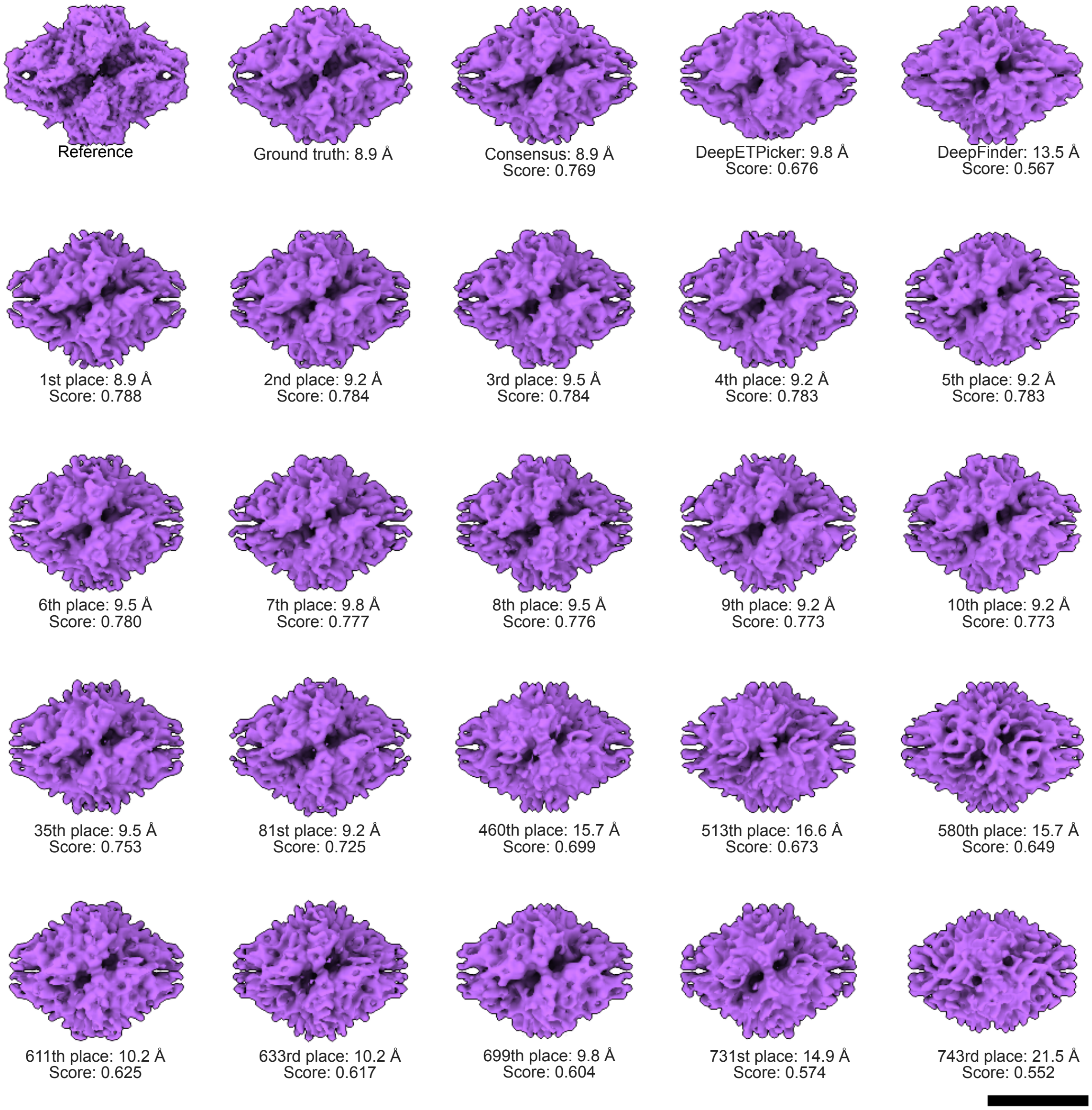


**Supplemental Fig. 2: Comparison of beta-galactosidase maps**. The scale bar corresponds to 100 Å.

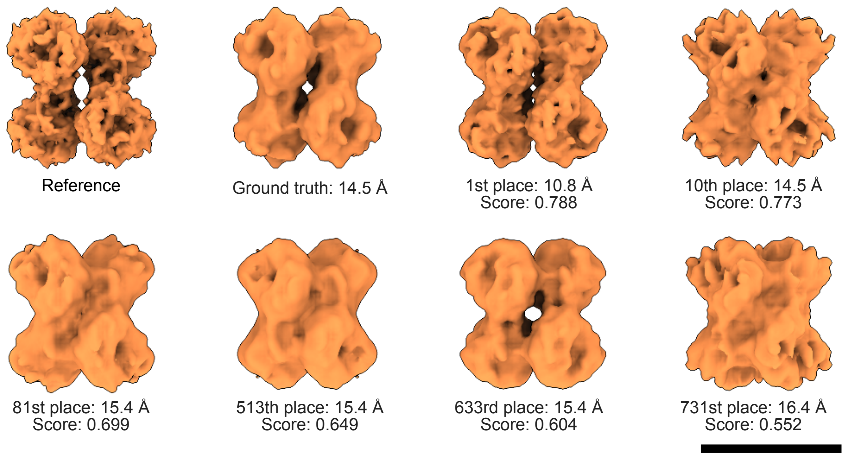


**Supplemental Fig. 3: Comparison of beta-amylase maps**. The scale bar corresponds to 100 Å.
