## Supplemental Note 2 for "Lessons learned from a Kaggle challenge for particle picking in cryo-electron tomography"

### Supplemental Note 2: Map quality assessment

In STA, gold standard refinement divides particles into half sets that are processed independently, with map resolution estimated from the Fourier shell correlation (FSC) between the refined half maps[36]. However, this half-maps FSC is especially prone to overestimating map quality for symmetric particles, since the application of symmetry can reinforce features that are consistent between half maps, even when those features are artifactual (Supplemental Fig. 4). An alternative approach to estimate resolution is map-model FSC[35], which compares the refined model to a reference map instead. Although the accuracy of this metric is limited if the reference map does not match the experimental map, for example due to compositional or conformational heterogeneity, we found that resolution estimates from map-model FSC better matched visual inspection and deteriorated as expected for maps dominated by symmetrization artifacts (Supplemental Fig. 4).

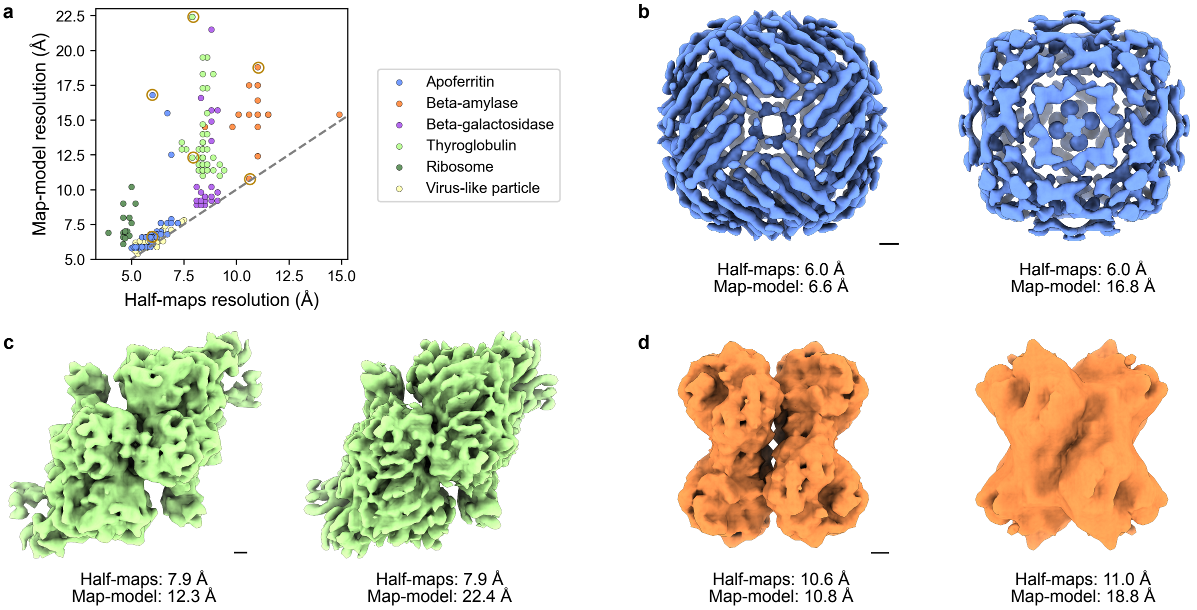


**Supplemental Fig. 4: Map-model FSC provides a more accurate resolution estimate than half-maps FSC, particularly for symmetric particles. a.** Resolution estimates from half-maps and map-model FSCs are compared for subtomogram averages for the indicated particle. The maps circled in yellow are visualized in panels **b-d**. Refined maps of **b.** apoferritin, **c.** thyroglobulin, and **d.** beta-amylase with comparable resolution estimates according to half-maps FSC and diverging resolution estimates based on the map-model FSC are visualized. The scale bar in each panel corresponds to 10 Å.

For beta-amylase, beta-galactosidase, and thyroglobulin, the reference model was generated by simulating density maps from atomic models deposited in the Protein Data Bank. This approach was also used to generate the ribosome reference map, though for this species a mask was applied to the small subunit to mitigate the impact of conformational heterogeneity — specifically, inter-subunit ratcheting — on the resolution estimate. For the apoferritin and virus-like particle assessments, the ground truth and contestants' maps were compared to experimental maps determined by single-particle cryoEM to ensure realistic density values for these proteins' shell-like interiors. Our simulation method treated this interior region as vacuum rather than solvent, which negatively impacted the resolution estimate (Supplemental Fig. 5). For these species, we additionally applied a mask to isolate the comparison to a single asymmetric unit (ASU), which further improved agreement with the reference map.

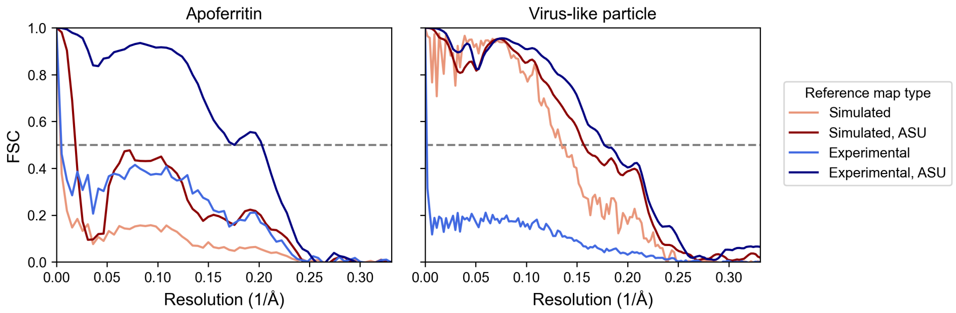


**Supplemental Fig. 5: The use of an experimental map as the reference improves map-model FSC agreement for shell-like particles.** Map-model FSC curves are compared when the reference map is either simulated from an atomic model in a way that treats the particle's unstructured interior as vacuum or an experimental map from single-particle cryoEM. A mask is applied so that only a single ASU is considered in the calculation for the indicated curves.

We also observed that resolution estimates sometimes varied between independent STA runs from the same starting coordinates. There are multiple sources of non-deterministic behavior during STA, such as random splitting of particles between half sets and particles' initial angular assignments. However, comparing independent STA runs for four of the six species showed that the difference in resolution between replicates was generally small for maps at subnanometer resolution (Supplemental Fig. 6a). Increased variability of >3 Å was observed for some replicates that yielded lower resolution maps but was less consequential since differences in structural features that are resolved at 5 versus 8 Å are much more significant than at 15 versus 18 Å. Though this variability makes it difficult to definitively determine which contestant's picks yielded the highest-quality map for each species, it does not change the interpretation of the results.

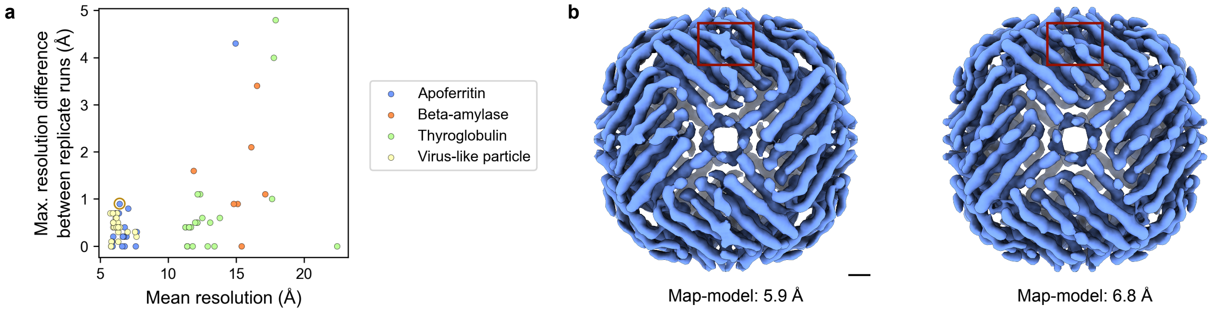


**Supplemental Fig. 6: Variability between STA replicates. a.** The maximum difference in resolution between replicate STA runs is plotted as a function of their mean resolution, with data points colored by species. Two maps from the replicate set circled in yellow are shown in panel b. **b.** Maps from the set of picks with the highest resolution difference between replicate runs are shown. Although certain regions like the one boxed in red showed observable differences in density continuity, the gross structure was largely the same despite the 0.9 Å difference in global resolution. The scale bar indicates 10 Å.
